# Selection navigates a degenerate circuit space: behavioral individuation without structural differentiation in constrained neuroevolution

**DOI:** 10.64898/2026.07.13.736323

**Authors:** Pranet Khetan, Aditya Asopa

## Abstract

Neural degeneracy, the capacity of structurally distinct circuits to perform the same function, is typically studied as an emergent property of biological neural systems. Here we impose it by construction, engineering the conditions that make degeneracy the expected outcome of evolutionary search, and ask what individual-target selection achieves within that degenerate space. We use constrained neuroevolution to evolve 14-neuron recurrent circuits (Dale’s Law, sparse connectivity, quantized weights) replicating the natural navigation behavior of 9 individual mice across 54 independent evolutionary runs. Architectural constraints impose a structural floor: no aggregate circuit statistic differs across mice (0/18 features, all *p*_FDR_ *>* 0.47), and this uniformity extends to the topology axis itself: topology distance predicts behavioral distance at no scale, in evolved or random constrained agents alike, and no structural axis carries significant information about behavioral identity (maximum NMI = 0.2173). Yet behavioral individuation is robust: own-mouse fitness error is 33.4% lower than cross-mouse error, and this specialization persists on held-out data. We show that what individual-target selection shapes is not circuit structure but the strength of functional sensitivity commitment: specialists develop roughly 6.4–6.7*×* higher sensitivity variance than generalists trained on all mice simultaneously (9/9 mice, mouse-level *p* = 0.002), despite exploring statistically indistinguishable topological diversity. The particular pathways a circuit commits to carry no shared mouse-specific signature, so behavioral individuation is expressed as a magnitude of functional commitment rather than a structural or pathway fingerprint—degeneracy that operates not just at the structural level but at the level of the computational strategies that implement individual behavior.

**Author summary:** Every mouse behaves a little differently from every other mouse. Where does that individuality live in the brain? The intuitive answer is that different behavior must come from differently wired circuits. We tested that assumption directly. We evolved small artificial neural circuits— just 14 neurons, built under biological rules such as Dale’s Law—until each one reproduced the natural exploratory behavior of one specific mouse, repeating the process 54 times across nine mice. The circuits captured their target well: each performed markedly better on the mouse it was evolved for than on any other, and this advantage held on behavior the circuits had never seen. Yet when we examined their wiring, we found nothing. No structural measurement we tried—connection statistics, network topology, representational geometry, or dynamics— could recover which mouse a circuit had been evolved to match. Individuality was robust in behavior and invisible in structure. What did differ was how strongly each circuit committed to particular sensory inputs, and even those pathways were idiosyncratic rather than shared. Individual differences can therefore be carried by the magnitude of functional commitment rather than by any structural signature.

## 1 Introduction

Neuronal circuit motifs are conserved across species and provide the structural substrate for a wide range of behaviors [Luo, 2021]. Yet individuals with nearly identical genomes can differ systematically in their navigation strategies [Rosenberg et al., 2021, Bogado Lopes et al., 2023]. Whether such behavioral individuality reflects differences in circuit structure, or whether a shared architecture can support diverse behavioral outputs through variation in its internal parameters, is a central unresolved question in systems neuroscience. This question is difficult to address experimentally: developmental programs constrain the phenotypic space available to selection [Tosches, 2017], and the scale and heterogeneity of biological circuits makes it hard to distinguish which structural features are required by behavior and which reflect developmental or phylogenetic history [Roberts et al., 2022]. Large-scale connectomics projects have produced near-complete wiring diagrams in several organisms [Consortium, 2024, MICrONS Consortium, 2025], yet the mapping from connection structure to circuit function remains a major unsolved problem [Bargmann and Marder, 2013, Jonas and Kording, 2017].

Here we take a computational approach. We use constrained neuroevolution to evolve minimal neural circuits that replicate the navigation behavior of individual mice, then ask what the solution space of these circuits reveals about the relationship between circuit structure and behavioral identity. Our model system is a 14-neuron recurrent network (6 sensory, 6 interneurons, 2 motor) navigating a simulated hierarchical maze, subject to three biological constraints: Dale’s Law [Strata and Harvey, 1999] (fixed excitatory/inhibitory identity, with all outgoing connections sharing the same sign), sparse connectivity (at most 3 incoming and 3 outgoing connections per neuron), and quantized synaptic weights (*{*0.25, 1.0*}*, reflecting the skewed distribution of cortical synaptic strengths in which a small fraction of connections operate at high efficacy [Song et al., 2005]). We evolve independent populations to replicate the natural, exploratory navigation behavior of 9 individual mice [Rosenberg et al., 2021], using a fitness function that tracks four behavioral statistics (transition probabilities, spatial occupancy, path tortuosity, and turn bias) rather than an external reward signal. This choice avoids presupposing what computations the circuit must implement and allows the fitness function to track each individual’s behavioral signature directly.

Neural degeneracy is the ability of structurally distinct elements to perform the same function [Edelman and Gally, 2001, Marder and Goaillard, 2006]. In this system degeneracy operates at two levels. With respect to *competence* (navigational fitness), the mapping is classically many-to-one: many structurally distinct circuits are each a sufficient substrate for the same function—being a fit navigator—so any circuit selection can reach within the constrained space navigates well. With respect to *identity* (which individual mouse a circuit was selected to replicate), structure is degenerate in a stronger sense: identity leaves no signature on any structural axis, even though the behaviors themselves differ. Individual specialisation is real but structurally invisible; it is carried functionally, not structurally. This identity-level regime is adjacent to—but distinct from—what Albantakis et al. [2024] term plurifunctionality (a single structure supporting *multiple distinct* functions): here each circuit implements one individual’s behavior, and what is degenerate is the mapping from structure to identity, not the number of functions per circuit. Degeneracy is a fundamental organizational principle of biological nervous systems. In naturally evolved circuits, degeneracy is difficult to study in isolation because structural variation, developmental noise, and evolutionary history co-vary. Our constrained system offers a different entry point: the architectural constraints (Dale’s Law, degree limits, quantized weights) impose degeneracy by construction, compressing all solutions into the same structural region regardless of the behavioral target. We can therefore treat the constrained circuit space as a controlled laboratory for degeneracy, and ask: what can degeneracy achieve under these conditions, and what are its limits?

Degeneracy in this system is pervasive and structurally invisible. Across 54 independent evolutionary runs, no aggregate circuit statistic differs across mice: a structural floor imposed by the constraints, not by convergent selection. This degeneracy extends beyond aggregate statistics. Connection topology varies freely within and across mice, yet topology distance predicts behavioral distance at no scale, and random constrained agents show the same flat relationship as evolved ones. No structural axis (topology, synaptic magnitude, or excitatory/inhibitory identity) carries more than weak, non-significant information about behavioral identity (maximum normalized mutual information = 0.2173 across all axes). Yet behavioral individuation is robust: own-mouse fitness error is 33.4% lower than cross-mouse error (specialization index = 0.334), and this specialization persists across all four behavioral dimensions and on held-out data. The question then is what topology is doing if it is not encoding behavioral identity. It is causally necessary: permuting connection targets disrupts motor output 3.53*×*, and this disruption is mouse-specific. But topological variation is degenerate with respect to behavioral identity; any topology drawn from the constrained space provides a sufficient computational substrate, and none is uniquely identifying. What individual-target selection shapes instead is the strength of functional sensitivity commitment: specialists develop roughly 6.4–6.7*×* higher sensitivity variance than generalists trained on all mice simultaneously, despite exploring statistically indistinguishable topological diversity. The particular pathways a circuit commits to carry no shared mouse-specific signature, so individuation is carried by the magnitude of commitment rather than a specific pathway signature. Selection navigates a degenerate space to produce individual-specific function without leaving a legible structural trace.

## 2 Results

### 2.1 Constrained neuroevolution reliably replicates individual mouse behavior

We evolved 54 independent populations (9 mice *×* 6 replicates, 500 agents each) for 150 generations. All runs converged reliably: population mean fitness changed by less than 2% over the final 30 generations in all 54 runs, with the steepest improvement in generations 1–50 and diminishing returns thereafter (Figure 1C–D; Supplementary Figures S3–S1). Per-mouse mean fitness at generation 150 ranged from 0.770 to 0.847 (grand mean 0.809 *±* 0.023; lower = better behavioral match), well below the random-agent baseline of 2.87–3.39. No run showed fitness divergence, premature convergence, or collapse.

**Figure 1.**
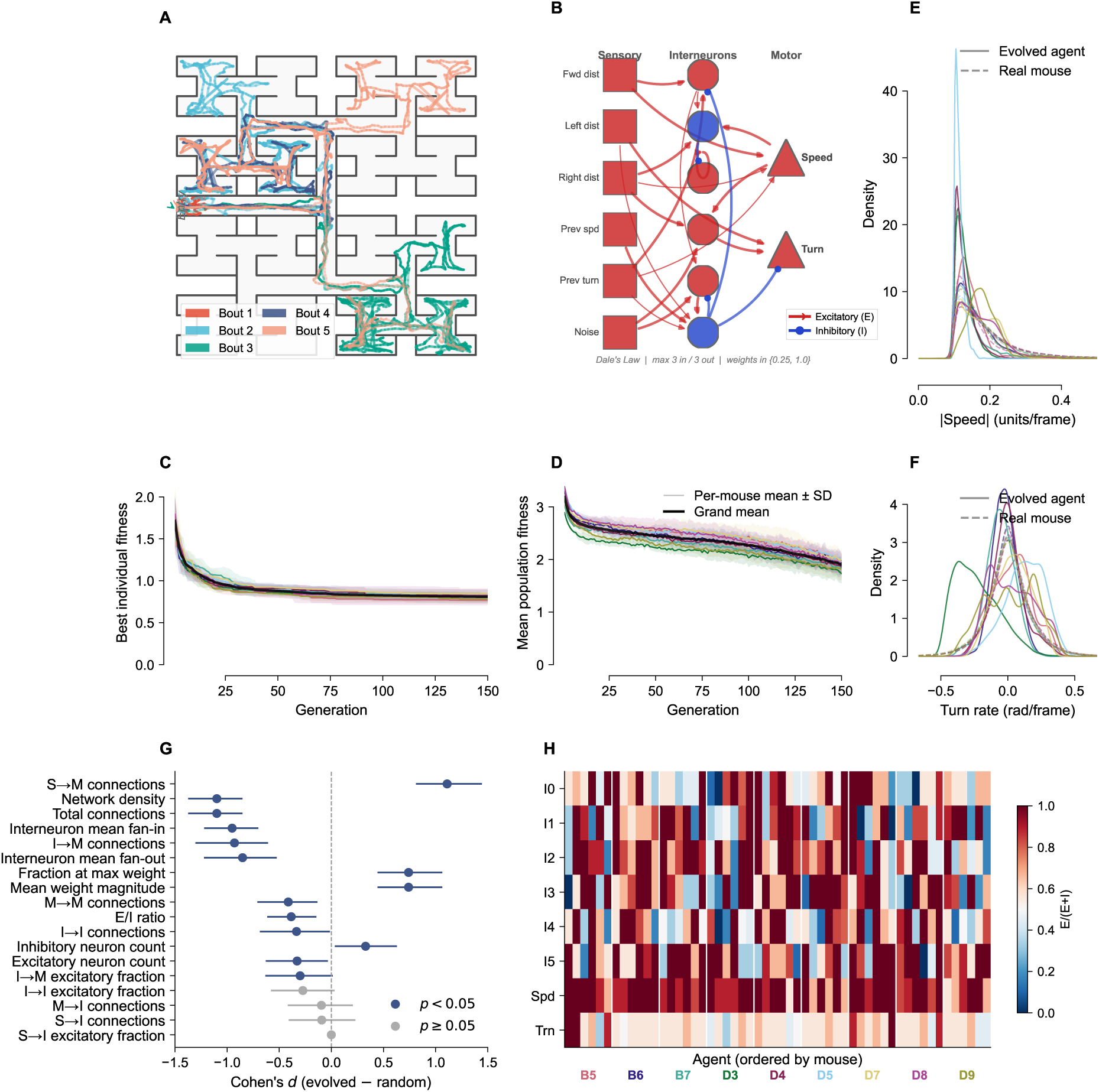
The constrained neuroevolution system. (A) Hierarchical binary Y-maze with representative mouse trajectories (5 bouts, B5). (B) 14-neuron recurrent agent architecture: 6 sensory inputs (squares), 6 interneurons (circles), 2 motor outputs (triangles). Excitatory connections in red, inhibitory in blue. Dale’s Law, sparse connectivity (max 3 in/3 out), and quantised weights 0.25, 1.0 are enforced throughout evolution. (C) Population mean fitness SD across 6 replicates per mouse over 150 generations (lower = better behavioral match). Fitness decreases steeply in generations 1–50, with slow improvement continuing to generation 100–120 before plateauing. (D) Best individual fitness per mouse. The best agent in each replicate plateaus earlier and more cleanly than the population mean, confirming that evolutionary improvement concentrates in the leading individuals. (E) Speed distributions for evolved agents (solid) and real mice (dashed), per mouse. Agents reproduce the characteristic speed profile of their target mouse without explicit selection for speed. (F) Turn rate distributions for evolved agents (solid) and real mice (dashed), per mouse. Agents reproduce the characteristic turn rate profile of their target mouse without explicit selection for turn rate statistics. (G) Evolved vs random circuit features: Cohen’s *d* with 95% bootstrap CI for all 18 aggregate structural features. Navy *p <* 0.05; grey *p* 0.05. Evolved circuits are sparser, more feedforward, and use stronger connections than random agents. (H) E/I balance at generation 150 for all 8 non-sensory neurons (54 8 heatmap, agents ordered by mouse; white lines separate mouse groups). Speed motor (Spd) converges uniformly to mean 0.85 excitatory across all mice; turn motor (Trn) converges to mean 0.64 with greater variability, consistent with closed-loop inhibitory modulation of turning.

Beyond the four fitness components, evolved agents reproduced behavioral properties not explicitly selected for (Supplementary Figure S4). Speed distributions showed the characteristic slow-fast pattern of real mice: slow exploratory phases interspersed with fast directed traversals (Figure 1E). The overall speed *scale* is partly set by the momentum constants (*α_v_*, *v*_max_, *ω*_max_), which are fixed from each mouse’s trajectory statistics (Methods); the emergent property is the temporal *patterning* of slow and fast phases, which the fitness function does not target directly. Evolved agents also reproduced the turn rate distributions of their target mouse (Figure 1F) and showed wall contact fractions consistent with thigmotaxis (Supplementary Figure S4), all without explicit selection for these properties. These results indicate that the evolved circuits capture a general behavioral policy rather than a narrow set of optimised statistics.

Evolution also selected a consistent circuit architecture across all 54 populations. Compared to randomly initialised constrained agents, evolved circuits are sparser, more feedforward, and use stronger individual connections: they carry more direct sensory-to-motor shortcuts, fewer recurrent pathways through interneurons, and a higher fraction of connections at the maximum weight magnitude (Figure 1G). Interneurons have reduced fan-in and fan-out, consistent with a supporting rather than central computational role. Representative network diagrams for all 9 mice are shown in Supplementary Figure S2.

Excitatory-inhibitory balance revealed a functional dissociation within this architecture. The speed motor neuron converged to a strongly excitatory-dominated regime (mean E/(E+I) *≈* 0.85) uniformly across all 9 mice, consistent with an open-loop tonic drive. The turn motor neuron converged to a more balanced ratio (mean *≈* 0.64) with greater variability between mice, consistent with closed-loop control in which inhibitory input modulates turning direction. Given that each motor is a single 2-input readout with E/(E+I) restricted to *{*0, ^1^ , ^2^ , 1*}*, we read this only as loose analogy to the push-pull organisation of biological locomotor circuits [Kiehn, 2006] (Figure 1H; Supplementary Figure S5 for the full temporal evolution). This dissociation is an emergent internal property of the evolved networks: it was not specified by the fitness function and reflects the constraint geometry rather than individual behavioral variation.

Together, these results establish that the evolutionary system works: all 54 runs produce agents that match their target mouse’s behavioral statistics, generalise to emergent behavioral properties, and converge on a shared feedforward-dominant architecture with a consistent E/I organisation. This shared architecture is the starting point for the degeneracy analysis that follows.

### 2.2 Architectural constraints impose structural degeneracy, yet enable behavioral individuation

Evolved circuits occupy a significantly narrower region of weight space than randomly initialised constrained agents (pairwise cosine similarity: evolved = 0.211 vs random = 0.070; Mann-Whitney *p* = 1.2 *×* 10*^−^*^229^; Supplementary Figure S6), but the resulting 54 *×* 54 similarity matrix shows no mouse-level block structure (Figure 2A). Evolution compresses all circuits toward a shared structural profile, not toward mouse-specific ones: this is structural degeneracy at the aggregate level.

**Figure 2.**
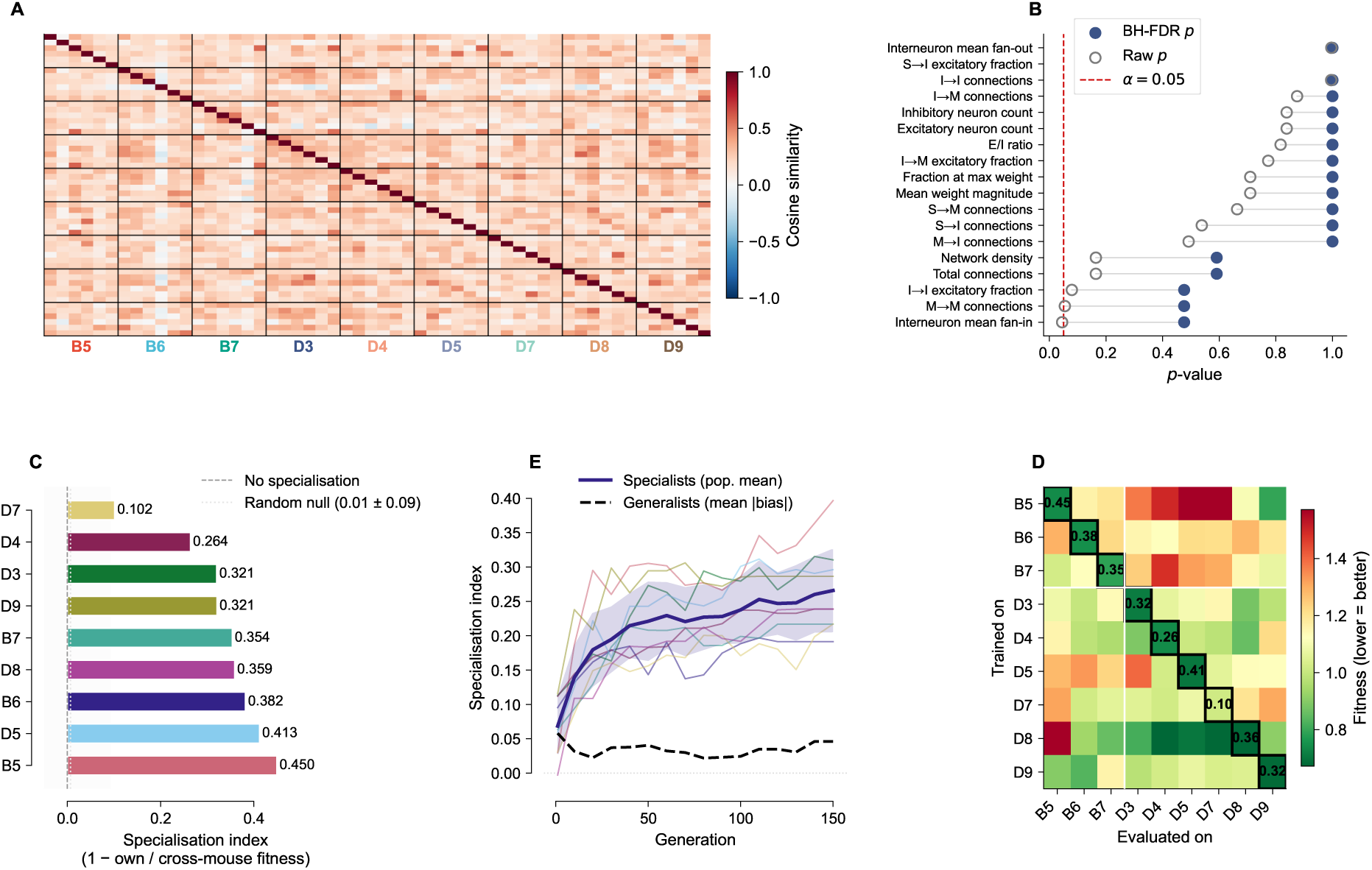
Architectural constraints impose structural degeneracy, yet behavioral individuation is robust. (A) 54 54 weight cosine similarity heatmap, agents ordered by mouse group (separators mark group boundaries). No mouse-level block structure is visible: evolution compresses all circuits toward a shared structural region, not toward mouse-specific clusters. (B) Raw (open circles) and BH-FDR-corrected (filled, navy) *p*-values for all 18 aggregate circuit features, sorted ascending by raw *p*, connected by grey lines. The smallest raw *p*-value is 0.045 (interneuron fan-in); after BH-FDR correction, all adjusted *p >* 0.47 (red dashed line: *α* = 0.05). Zero of 18 features differ across mice regardless of correction method. (C) Per-mouse behavioural specialisation index (1 own/cross fitness), sorted descending. All 9 mice show positive indices (dashed line = no specialisation). (D) 9 9 cross-mouse generalisation matrix. Colour encodes fitness (lower = better match), diverging palette centred on the off-diagonal mean. Bold boxes highlight diagonal (own-mouse) entries; diagonal annotations show per-mouse specialisation index. (E) Specialisation index over 150 generations: coloured per-mouse traces with bold population mean (thick line; grey band = 1 SD). Dashed black line shows generalist mean bias , which remains near zero throughout. Specialisation is absent at generation 1 and emerges exclusively under individual-target selection pressure.

Testing 18 aggregate structural features with one-way ANOVA (Benjamini-Hochberg FDR; Benjamini and Hochberg 1995) yields 0/18 significant results. The smallest raw *p*-value is 0.045 (interneuron fan-in); after BH-FDR correction, all adjusted *p >* 0.47 (Figure 2B). This null result reflects an architectural floor rather than an evolved property: the same ANOVA, run on an equal-sized matched sample of randomly initialised (unevolved) constrained agents (54 agents in a matched 9-group *×* 6 design, identical power to the evolved test), returns an identical 0/18. The degree limit, quantized weights, and Dale’s Law constrain the space of realisable circuits so tightly that no aggregate structural feature can diverge across mice regardless of training target. The study is adequately powered to detect the observed effect sizes: the three largest-effect features each exceed the 80% power threshold, and their non-significance reflects the absence of a detectable effect (Supplementary Figure S7). Both magnitude features likewise show no between-mouse differences (*p*_FDR_ = 1.00). Every measurable aggregate structural property converges to a common floor.

Despite this structural uniformity, evolved agents are strongly specialised to their training mouse. We evaluated each mouse’s best agent on all 9 behavioral targets to construct a 9 *×* 9 cross-mouse generalisation matrix (Figure 2D). Within-mouse fitness averaged 0.748 *±* 0.113 (diagonal; lower = better match), compared to 1.122 *±* 0.242 for cross-mouse evaluations (off-diagonal). The mean behavioral specialisation index (1 *−* own/cross fitness) was 0.334: own-mouse fitness error is 33.4% lower than cross-mouse error. All 9 mice showed positive indices (B5 most specialised, index = 0.450; D7 least, index = 0.102; Figure 2C). Specialisation is consistent across all four fitness components (Supplementary Figures S9–S10) and within each genetic strain independently of any strain-level grouping (Supplementary Figures S11 and S12; Supplementary Table S2). Randomly initialised constrained circuits show specialisation indices near zero (mean 0.007 *±* 0.086; Figure 2C), confirming that individual-target optimisation (not the constrained architecture itself) produces specialisation. Specialisation persists on heldout trajectory data (Supplementary Analysis 3.19). Specialisation emerges progressively with individual-target training: the population mean specialisation index rises from 0.068 at generation 1 to 0.266 at generation 150 (Figure 2E), while generalist agents trained simultaneously on all nine mice remain near zero throughout (mean *|*bias*|* at generation 150: 0.046). Per-mouse trajectories are shown in Supplementary Figure S22.

Circuits with identical aggregate structure produce specialised behavior. The constraint geometry compresses structural variation by design; behavioral individuation is robust. The remainder of this paper characterises how this is possible: what degeneracy achieves in this space, and what selection must do within it.

### 2.3 Degeneracy extends to the topology axis: topology variation does not predict behavioral identity

The structural floor established in Section 2.2 operates at the level of aggregate statistics. Does degeneracy also extend to the specific pattern of connections? If topology were the axis on which mice differ, we would expect agents with similar topologies to produce similar behavioral profiles, and agents trained on the same mouse to converge to more similar topologies than agents trained on different mice. Neither is the case.

We computed pairwise Spearman correlations between topology distance (Jaccard) and behavioral profile distance (cosine) across all 1431 agent pairs (Figure 3B–C). The overall correlation was *ρ* = 0.001 (*p* = 0.966), indistinguishable from zero. Splitting by pair type makes no difference: within-mouse pairs (agents trained on the same mouse) give *ρ* = *−*0.026 (*p* = 0.765); between-mouse pairs give *ρ* = 0.008 (*p* = 0.764). Agents with very similar topologies are no more behaviorally similar than agents with very different topologies. Nor did individual-target selection converge topology within mouse groups: within-mouse Jaccard distances (mean = 0.843) are indistinguishable from between-mouse (mean = 0.840; Mann-Whitney *p* = 0.493; Figure 3B). The 9 *×* 9 behavioral specialization structure is robust (Mann-Whitney *p* =*<* 0.001 for within vs between behavioral distance), but it is invisible in the topology distance matrix.

**Figure 3.**
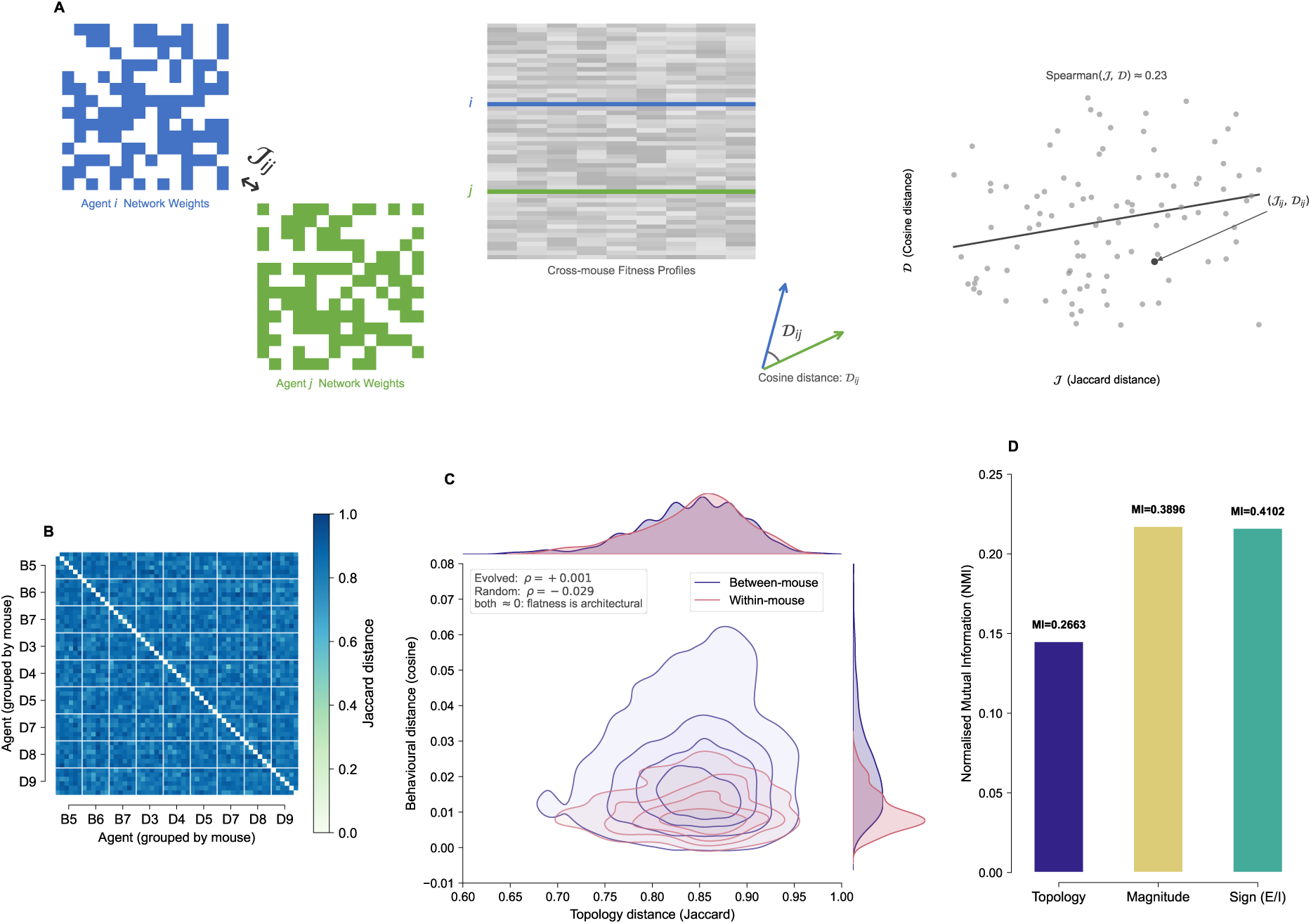
Structural degeneracy extends across all measurable axes. (A) Analysis schematic: each agent is characterised by a binary 14 × 14 topology matrix and a 9-dimensional behavioural profile vector; pairwise Jaccard distance *J* (topology) and cosine distance (behaviour) are computed across all 1431 agent pairs, and Spearman *ρ* quantifies their correlation. (B) 54 × 54 topology Jaccard distance heatmap (agents ordered by mouse group; white lines mark boundaries). No within-mouse block structure is visible: within-mouse topology distances (mean = 0.843) are indistinguishable from between-mouse (mean = 0.840; MW *p* = 0.493). Contrast with the behavioural specialisation block structure in Figure 2A. (C) Joint KDE of topology Jaccard distance vs behavioural cosine distance for all 1431 agent pairs (contours only; rose = within-mouse, indigo = between-mouse). Top marginal: topology distance KDE — within-and between-mouse distributions overlap completely (MW *p* = 0.493), confirming topology flatness. Right marginal: behavioural cosine KDE — within-mouse distances are lower than between-mouse (MW *p* =*<* 0.001), confirming specialisation. Spearman *ρ* annotated for both evolved (*ρ* = 0.001, *p* = 0.966) and randomly initialised constrained agents (*ρ* = −0.029, *p* = 0.270); both are indistinguishable from zero: topology-behaviour flatness is architectural. (D) Normalised mutual information (NMI) between each structural axis and mouse identity, computed via *k*-means clustering (*k* = 5) across all 54 agents. Maximum NMI = 0.2173 across all three axes; all at or below chance expectation (adjusted MI near zero or negative across *k* = 3–9). Reference line at NMI = 0.5 provides a calibration anchor for moderate association.

The same null holds within each mouse. Among the 135 within-mouse replicate pairs, neither topology distance (*ρ* = 0.034, *p* = 0.696) nor magnitude distance (*ρ* = 0.045, *p* = 0.601) predicts fitness difference. Partial correlations controlling for the other axis are equally null (topology controlling for magnitude: *ρ* = *−*0.018, *p* = 0.835). The within-mouse fitness landscape is many-to-one on both structural axes: many topologies achieve equivalent fitness for the same behavioral target.

Critically, this flatness is not a product of selection. We evaluated 54 randomly initialised constrained agents (same architecture, no training) on all 9 behavioral targets and computed the same topology-behavior correlation. Random agents show *ρ* = *−*0.029 (*p* = 0.270), and their within/between behavioral distance structure is absent (Mann-Whitney *p* = 0.998; Figure 3C). Degeneracy on the topology axis is thus architectural: the constraint geometry prevents topology from encoding behavioral identity.

### 2.4 No structural axis carries behavioral identity information

The correlational analyses of Section 2.3 establish that topology does not predict behavioral identity. To quantify degeneracy across all three structural axes (connection topology, synaptic magnitude, and excitatory/inhibitory sign) simultaneously, we computed mutual information (MI) between each axis and behavioral identity (the 9-category label indicating which of the 9 mice each agent was trained on), following the framework of Tononi et al. [1999].

For each structural axis, we represented all 54 agents as feature vectors and assigned cluster membership via k-means (*k* = 5), grouping agents by structural similarity on that axis. MI and normalised MI (NMI) were then computed between cluster assignment and mouse identity (Figure 3D; Methods, Section 3.16; Supplementary Figure S15). No axis carries more than weak information: NMI = 0.1451 for topology, 0.2173 for magnitude, and 0.2161 for excitatory/inhibitory sign. The maximum NMI across all axes is 0.2173, well below NMI = 0.5 (moderate association; reference line, Figure 3D), and at or below chance expectation (adjusted MI near zero or negative across *k* = 3–9; Methods, Section 3.16). The system is degenerate on every measurable structural dimension simultaneously.

### 2.5 Connection topology is causally necessary but structurally degenerate; no structural decomposition fingerprints identity

The analyses above establish that topology variation is uninformative about behavioral identity. This raises a sharp question: is topology doing anything at all? The answer is yes: topology is causally necessary, but its role is causal rather than informational. These are distinct questions with distinct evidential structures. The permutation analysis is a causal intervention: it asks whether *this* agent’s specific topology is necessary for *its* computation. The correlation analyses of Sections 2.3–2.4 are informational: they ask whether topology variation *across independently evolved agents* encodes behavioral identity. That topology is causally necessary for each individual circuit is fully consistent with topology variation being informationally flat across circuits: any topology drawn from the constrained space provides a sufficient causal substrate, but no specific topology is uniquely identifying.

#### Topology is causally necessary for computation

We performed a source-preserving topology permutation: the specific source-to-target identity of each connection (which neuron wires to which) was scrambled while every aggregate structural feature (pathway counts, weight magnitudes, E/I composition) was preserved exactly. The manipulation therefore isolates specific wiring identity from the 18 aggregate statistics. Across all 54 agents, permutation elevated motor output MSE by 3.53*×* relative to replicate pairs (*p* = 8.6 *×* 10*^−^*^7^, Mann-Whitney; Figure 4B). This disruption is mouse-specific: permuted agents lost significantly more behavioral fitness on their own training mouse than on other mice (Δfit_own_ = 2.530 vs Δfit_other_ = 2.088; Wilcoxon signed-rank *p* = 0.0039, unanimous across all 9 mice; Figure 4A). A suggestive positive association links the degree of specificity to behavioural specialisation: the per-mouse specialisation index trends positively with the own/other Δfit ratio (Spearman *ρ* = 0.68, permutation *p* = 0.050, 95% bootstrap CI = [*−*0.10, 1.00], *n* = 9; Figure 4C). The wide bootstrap interval at *n* = 9 means this dose-response relationship is indicative rather than firmly established, and does not bear the weight of the causal conclusion, which rests on the within-agent permutation effect above. The specific pattern of connections matters causally for each agent’s computation.

**Figure 4.**
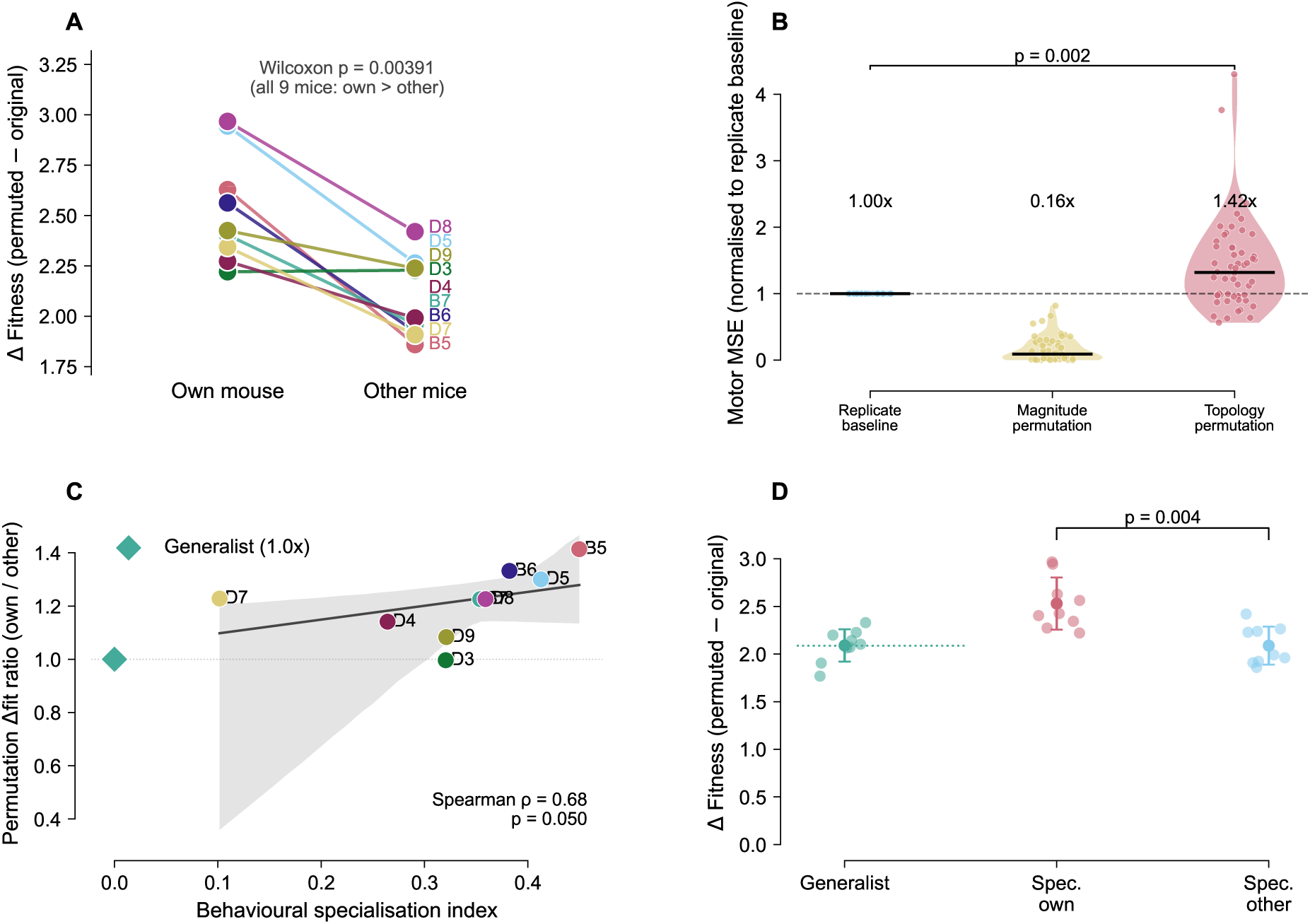
Topology is causally necessary but structurally degenerate. (A) Slope graph: permutation Δfit on own mouse vs other mice for each of the 9 specialist mice (coloured lines). All 9 lines slope upward; own-mouse degradation is unanimously greater (Wilcoxon *p* = 0.0039). (B) Motor MSE under topology permutation, magnitude permutation, and replicate baseline. Violin + strip plots; median marked. Topology permutation elevates MSE (1.419 baseline); magnitude permutation reduces it (0.164), reflecting architectural degeneracy in weight strengths. (C) Dose-response: permouse behavioural specialisation index vs permutation Δfit ratio (own/other). More specialised mice show greater permutation specificity (Spearman *ρ* = 0.68, permutation *p* = 0.050, 95% bootstrap CI = [ 0.10, 1.00], *n* = 9). The generalist (teal diamond) anchors ratio = 1.00 . (D) Generalist control: permutation Δfit for generalist agents, specialist-own, and specialist-other conditions (mean SD, individual points overlaid). Generalists carry no own-mouse peak: their mean Δfit (2.090, 15 replicates) sits at the specialist *other*-mouse level (2.088), well below the specialist *own*-mouse level (2.530), whereas specialists show unanimous own *>* other (Wilcoxon *p* = 0.0039). Generalist Δfit does vary across the nine test mice (Kruskal-Wallis *p* = 1.6 10*^−^*^10^), but this reflects intrinsic between-mouse disruptability shared identically across replicates (Friedman *p* = 9.3 10*^−^*^14^ for a consistent per-mouse rank order), not a per-individual calibration signature. Per-source ablation (54 agents 14 source neurons) is shown in Supplementary Figure S14.

#### Topology, not magnitude, is the computational substrate

To distinguish the role of connection targets from connection strengths, we applied a magnitude-only permutation: the *{*0.25, 1.0*}* strength assignments were shuffled among each source neuron’s existing connections while targets were held fixed. This produced an MSE ratio of 0.164*×*, *below* the replicate baseline of 1.0*×* (Table 1). Shuffling weight strengths makes agents *more* similar in motor output, not less: with at most 3 connections per source and only two magnitude values, 61.9% of source neurons carry uniform magnitude assignments and are unaffected by magnitude shuffling. Synaptic strength is a degenerate causal axis; connection topology is not.

**Table 1.** Permutation MSE metrics (normalised to replicate baseline). *p*-values are Mann-Whitney vs replicate pairs.

| Condition | MSE ratio | $p$ |
| --- | --- | --- |
| Topology permutation | $1.419\times$ | $8.6 \times 10^{-7}$ |
| Magnitude permutation | $0.164\times$ | 1.0000 |
| Between conditions | $p = 1.8 \times 10^{-302}$ | |

#### No structural decomposition fingerprints mouse identity

If the distributed topology pattern is causally necessary, does any structural component (local or distributed) carry mouse-specific information? We tested this at two levels. First, per-source ablation: permuting each neuron’s outgoing connections individually yielded a 54 *×* 14 sensitivity matrix with no mouse-level organisation: 0 of 14 source neurons significant after Bonferroni correction, and within-mouse sensitivity profiles no more correlated than between-mouse profiles (*r*_within_ = 0.150, *r*_between_ = 0.167, Mann-Whitney *p* = 0.6919; Supplementary Figure S14). Second, even if no single neuron is a fingerprint, the combined *pattern* of sensitivities across all neurons might be mouse-specific. We tested this with a representational similarity analysis (RSA): within-mouse agent pairs were no closer in 14-dimensional sensitivity space than between-mouse pairs (Mann-Whitney *p* = 0.529), and a Mantel test between the 54 *×* 54 sensitivity similarity matrix and a mouse-identity matrix found no correlation (Mantel *ρ* = *−*0.044, permutation *p* = 0.357; Supplementary Figure S16).

Connection topology is causally necessary for computation and is individually calibrated to each mouse’s behavioral target. But the specific topology an agent uses is degenerate with respect to behavioral identity: agents with very different topologies can achieve the same behavioral fitness, and no structural decomposition (individual neurons or the distributed pattern of ablation sensitivities) identifies a mouse from its circuit. What individual-target selection achieves within this degenerate structural space is the question the following section addresses.

### 2.6 Selection shapes functional sensitivity commitment; individual-target training is necessary

Having established that degeneracy operates on every structural axis, we ask what individual-target selection actually achieves. We trained fifteen generalist agents (independent replicates) simultaneously on all 9 mice, using fitness equal to the mean across all 9 behavioral targets, with the same architecture, constraints, and evolutionary parameters as the per-mouse runs.

#### Generalists and specialists share statistically indistinguishable topological diversity

Generalist agents converged structurally, as expected from the architectural floor (Section 2.2). Pairwise topology cosine similarity was 0.261 for generalists and 0.268 for specialists (Mann-Whitney *p* = 0.173; Figure 5A). Individual-target training does not produce a more committed or distinctive topology. The axis on which selection acts is not topological.

**Figure 5.**
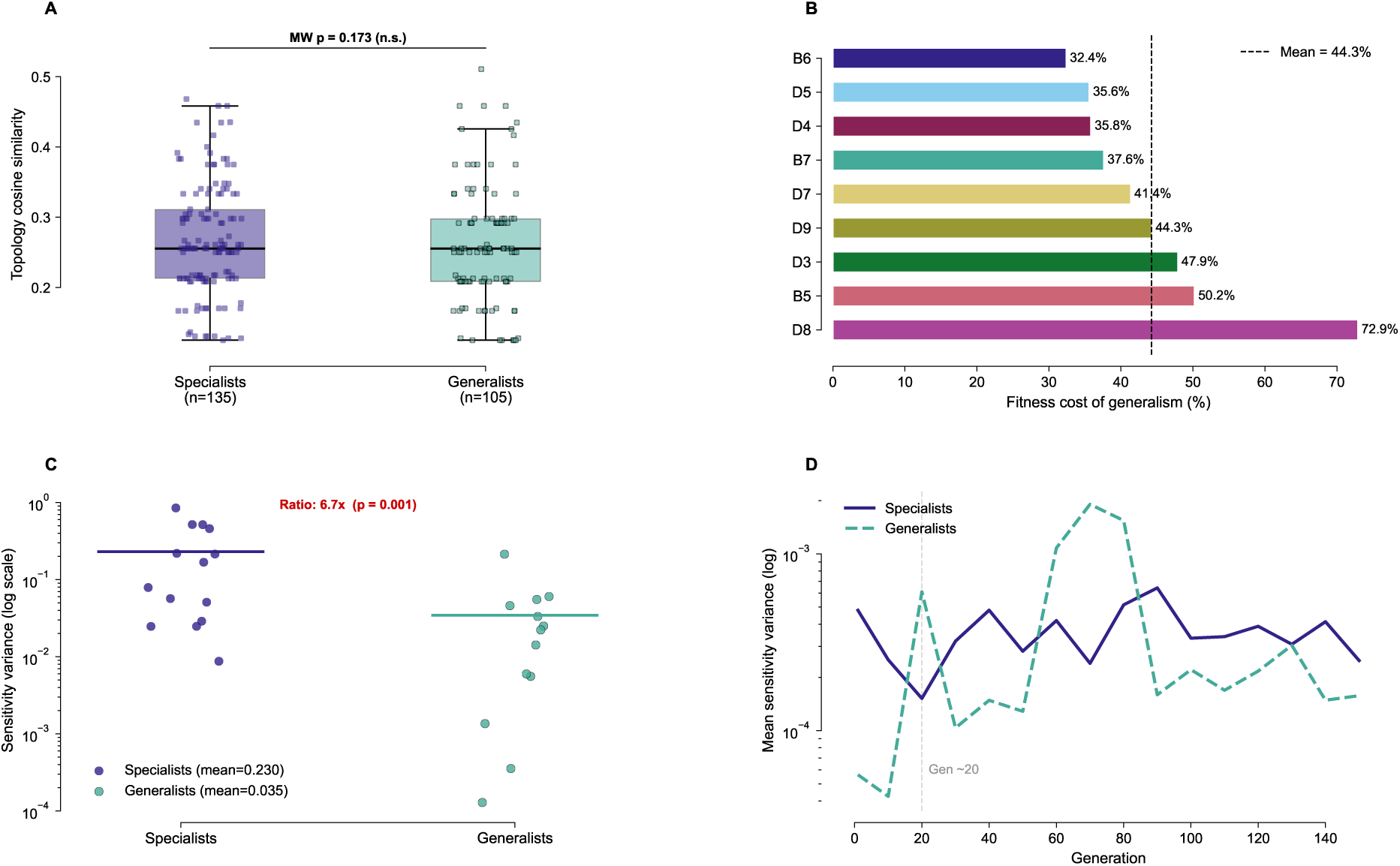
Selection shapes functional sensitivity commitment. (A) Topology cosine similarity for generalist and specialist agents (box + strip, Mann-Whitney *p* = 0.173). Generalists and specialists show indistinguishable topological diversity, confirming that individual-target training does not act on topology. (B) Fitness cost of generalism per mouse (percentage worse than specialist fitness), sorted descending. Dashed line = mean (44.3%). (C) Sensitivity variance per neuron for specialists vs generalists (log scale; individual neuron points shown, *n* = 14 per group). Specialists show high, varied sensitivity (*σ̄* ^2^ = 0.230); generalists show substantially lower sensitivity variance (*σ̄* ^2^ = 0.035). The specialist:generalist variance ratio is 6.4–6.7 (within-mouse-pooled to across-all-54 definitions; hierarchical-bootstrap 95% CI [2.88, 11.92]). At the mouse level (the independent unit), 9/9 mice exceed the generalist (one-sample Wilcoxon *p* = 0.002); the per-neuron Mann-Whitney is *p* = 0.001 (one-sided). (D) Mean sensitivity variance over 150 generations: specialists (solid, blue) vs generalists (dashed, orange), log scale. The endpoint shown in Panel C is the product of a progressive process: specialist sensitivity variance diverges from the generalist level from approximately generation 20 onward, and continues to rise throughout training. Individual-target selection specifically builds functional commitment; generalist training does not.

#### Individual-target training shapes functional sensitivity commitment

What it does produce is a committed functional sensitivity structure. We measured sensitivity variance per neuron (the variance of each neuron’s ablation sensitivity across replicates) as a readout of how consistently a circuit commits to particular pathways. Specialist agents show high, varied sensitivity (*σ̄* ^2^ = 0.230 per neuron): individual-target training produces circuits that are each strongly committed to perturbations of specific pathways, but which pathways vary across replicates, even within the same mouse. Consistent with the RSA null above (Supplementary Figure S16), within-mouse sensitivity profiles are no more similar to each other than to profiles from a different mouse: at generation 150 the pairwise cosine similarity is if anything marginally lower within-mouse (0.20) than between-mouse (0.24), but this small gap does not survive a pseudoreplication-aware test (label-permutation *p* = 0.076; the naive Mann-Whitney *p* = 0.040 treats the non-independent agent-pairs as independent and is anticonservative). We therefore do not claim that replicates commit to *distinct* pathways; the defensible reading is that no shared, mouse-specific pathway signature emerges. Selection builds the strength of functional commitment, not convergence to specific pathways. Generalist agents show substantially lower sensitivity variance (*σ̄* ^2^ = 0.035; ratio = 6.4–6.7*×*, within-mouse-pooled to across-all-54 definitions; per-neuron Mann-Whitney *p* = 0.001, one-sided, *n* = 14 neurons per group; Figure 5C), reflecting weaker, more distributed sensitivity to perturbations across all pathways. Because the ratio’s magnitude depends on the generalist replicate sample, we estimated it over fifteen independent generalist replicates and quantified its uncertainty with a hierarchical bootstrap (95% CI [2.88, 11.92]); treating the mouse as the independent unit, all 9 of 9 mice exceed the generalist (one-sample Wilcoxon *p* = 0.002), so the direction is unambiguous even though the point magnitude carries wide intervals. Generalists achieve comparable structural diversity at the cost of functional specialization of any kind.

#### Individual targets are necessary: the fitness cost of generalism

Generalist behavioral fitness was 44.3% worse than specialist fitness on average across mice (range 32.4–72.9%; Figure 5B). The hardest mouse to generalise across was D8 (72.9% cost) and the easiest was B6 (32.4% cost). The generalist’s permutation profile confirms the mechanism: its mean Δfit (2.090, 15 replicates) sits at the specialist *other*-mouse level (2.088)—not the specialist *own*-mouse level (2.530)—so the generalist carries no own-mouse peak, in contrast to specialists whose own-mouse degradation is unanimous (own *>* other in 8/9 mice, Wilcoxon *p* = 0.0039; Figure 4D). Generalist Δfit does differ across the nine test mice (Kruskal-Wallis *p* = 1.6 *×* 10*^−^*^10^), but this is an intrinsic between-mouse disruptability effect—the same mice are hardest to perturb in every replicate (Friedman *p* = 9.3 *×* 10*^−^*^14^)—rather than a per-individual calibration signature. A generalist circuit is thus disrupted like a specialist evaluated on its wrong mouse: partially calibrated to everyone, fully calibrated to none.

The fitness cost confirms that generalists do not simply fail to converge: relative to the per-mouse specialist fitness (mean of replicates), generalist error is higher by 43.6% on a ratio-of-means basis (per-mouse range 32.4–72.9%), yet they still reproduce each mouse’s behaviour well above chance, confirming clear behavioral learning. The low sensitivity variance is therefore not underfitting: it reflects the fact that generalist replicates, each having learned a working solution, show weaker and more distributed sensitivity to pathway perturbations, with no strong commitment to any particular subset of pathways.

The A4 sensitivity RSA result (Supplementary Figure S16) and the A6 sensitivity variance result are compatible and jointly clarify the mechanism. The *geometry* of the sensitivity profile does not fingerprint mouse identity (A4: which neurons an agent is sensitive to carries no consistent mouse-specific signature). What individuates a specialist is the *strength* of its sensitivity commitment: how strongly, and how variably across replicates, its circuits commit to specific pathway perturbations, with no shared mouse-specific pattern in which pathways are committed to. Individual-target training builds this commitment; generalist training cannot (Figure 5D). This commitment co-develops with behavioural specialisation throughout training: sensitivity variance in specialist circuits remains elevated above the generalist level from approximately generation 20 onward (Supplementary Figure S23); the small within-versus between-mouse gap in sensitivity-profile similarity does not survive a pseudoreplication-aware test (§2.6), so it is not evidence that replicates commit to distinct pathways.

What individual-target selection shapes is therefore not a distinctive topology, but the *strength* of a circuit’s functional sensitivity commitment: specialists develop higher sensitivity variance across replicates than generalists, while the RSA null (Supplementary Figure S16) shows that the specific committed pathways carry no consistent mouse-specific signature. Behavioral individuation in this constrained system is thus expressed as a magnitude of functional commitment, not as a distinctive circuit structure or a mouse-specific pathway signature.

### 2.7 Representational geometry is also degenerate

The structural, causal, and functional analyses above establish degeneracy across multiple levels of circuit description. Does this degeneracy extend to the representational level (to the geometry of neuronal activity embeddings)? A theoretical framework by Pezon et al. [2026] makes this question precise: even when different circuits produce identical latent dynamics, circuit structure may leave a “footprint” in neuronal embedding geometry, defined as the row loadings of the activity matrix under principal components analysis (PCA). The framework holds that circuit structure constrains the distribution of single-neuron functional properties even when the low-dimensional dynamics are equivalent across circuits. We operationalise this as a concrete, testable prediction about embedding geometry: if circuit structure leaves such a footprint, circuits with distinct architectures should differ in the intrinsic dimensionality or connected-component structure of their neuronal embeddings, even when their behaviorally relevant dynamics are identical.

We tested this prediction directly. For each of the 54 best-evolved agents (one per mouse per replicate), we re-simulated navigation across 20 bouts of up to 2000 frames each and recorded the full 14-dimensional neural state at every timestep. PCA was applied to the resulting (*N*_neurons_ *× T*_total_) activity matrix for each agent, retaining the top 6 components (which explained >90% of variance in most agents; Supplementary Figure S17), yielding neuronal embedding coordinates of shape 14 *×* 6. Before testing the main question, we verified that these embeddings are informative at the small scale of our system (*N* = 14): the two motor neurons (speed, neuron 12; turn, neuron 13) were consistently more separated from each other in PC1–PC2 loading space than the typical interneuron pair (0.583 *±* 0.220 versus 0.466 *±* 0.080; Mann-Whitney *p <* 0.001; Supplementary Figure S17). Note that PC loading orientation varies across agents due to the sign ambiguity of independent PCA solutions; all pairwise comparisons use Procrustes alignment to remove this ambiguity.

The manifold dimensionality of these circuits was near-maximal: the effective dimensionality at the 90% variance threshold was 5.83 *±* 0.37 out of a maximum of 6 retained components, and the participation ratio was 4.44 *±* 0.43, with no mouse-specific pattern (Supplementary Figure S18). Individual-target circuits do not converge on a low-dimensional representational cluster; they explore almost the full embedding space available to them.

To test whether individual-target selection produces convergence in activity embedding space, we computed pairwise Procrustes distances between the PC loading matrices of all 54 *×* 53/2 = 1,431 agent pairs, using orthogonal Procrustes alignment to remove the sign and rotation ambiguity introduced by independent PCA solutions. Within-mouse pairs (*n* = 135; 15 pairs per mouse across 6 replicates) showed Procrustes distances of 1.974 *±* 0.151, statistically indistinguishable from between-mouse pairs (*n* = 1,296; 1.988 *±* 0.160; Mann-Whitney *p* = 0.260; Figure 6A). This null result was robust across six independent similarity metrics spanning three activity-matrix construction modes (Procrustes; *p* = 0.14–0.26), representational dissimilarity analysis (*p* = 0.274), centred kernel alignment (*p* = 0.625), and covariance Frobenius distance (*p* = 0.745; Supplementary Figure S19). Generalist agents, which were not trained on any individual mouse, occupied the same region of embedding space as specialists (1.946 *±* 0.176; Mann-Whitney versus specialist within-mouse pairs, *p* = 0.920; Figure 6A).

**Figure 6.**
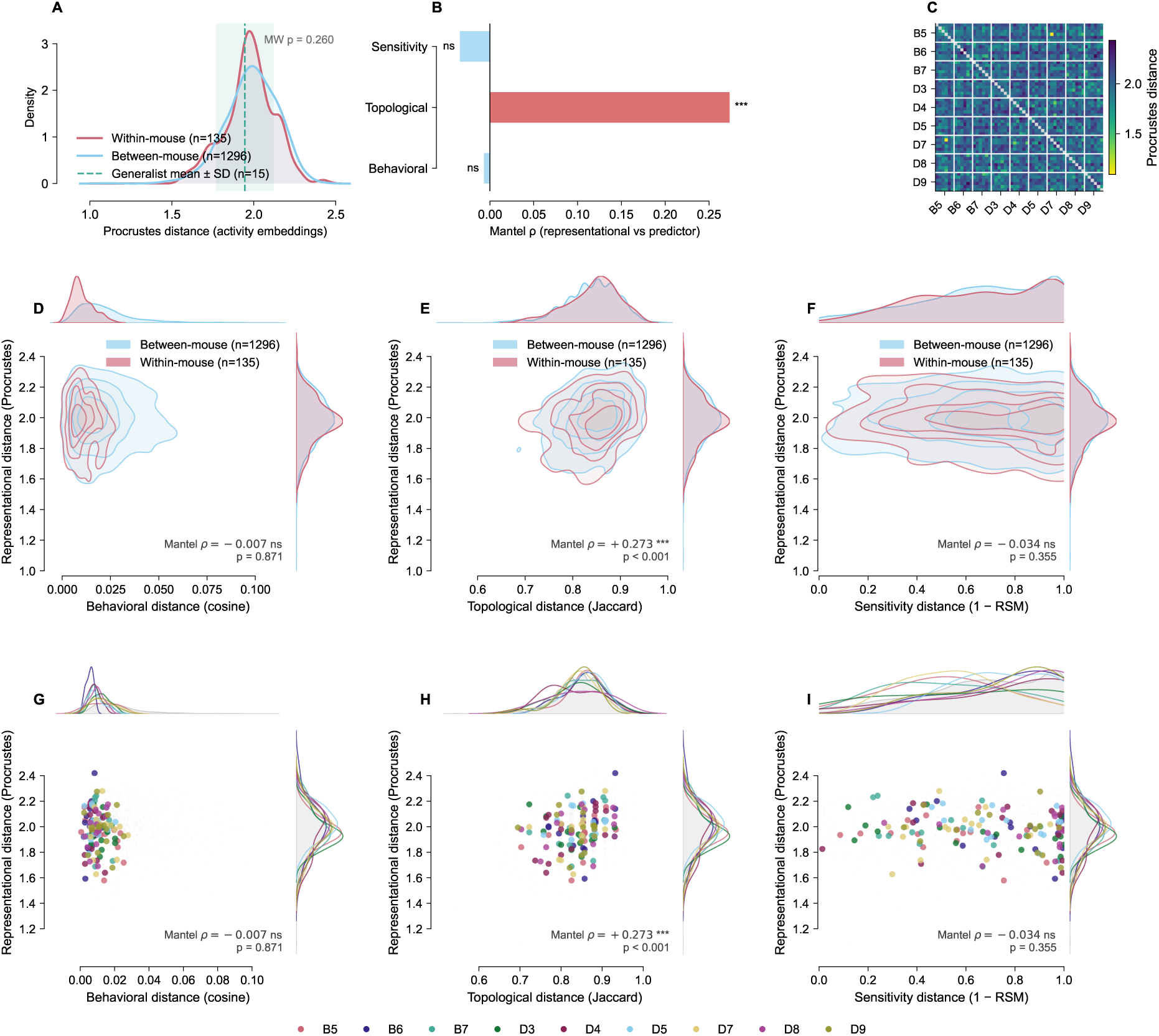
Representational geometry is degenerate across levels. (A) Procrustes distance between PC loading matrices (activity embeddings), within-mouse pairs (red, *n* = 135) versus between-mouse pairs (blue, *n* = 1,296). Dashed line and shaded band show generalist mean SD (*n* = 15). Distributions are indistinguishable (MW *p* = 0.260), confirming that individual-target selection produces no convergence in embedding space. (B) Mantel *ρ* between representational distance and three candidate predictor distances (behavioral, topological, sensitivity). Red bars indicate significant results; only topological distance is significant (*ρ* = 0.273, *p <* 0.001, _***_), in the expected *positive* direction: circuits sharing more topological scaffold tend toward slightly more similar representations. Behavioral and sensitivity correlations are null (*ρ* = 0.007 and *ρ* = 0.034, both *p >* 0.3, ns). (C) 54 54 Procrustes distance matrix, agents sorted by mouse identity (white lines mark 6 6 group boundaries). Absence of darker blocks on the diagonal confirms no within-mouse convergence in representational space. (D– F) KDE joint plots of representational distance versus behavioral (D), topological (E), and sensitivity (F) distances for all 1,431 agent pairs. Within-mouse (red) and between-mouse (blue) contours overlap on all three axes; Panel E reveals the only significant relationship (*ρ* = 0.273, *p <* 0.001). (G–I) Same three axes as D–F, with between-mouse pairs shown in grey and within-mouse pairs coloured by individual mouse identity (legend below). No mouse’s replicates cluster at distinctively lower representational distances, confirming that representational degeneracy holds at the level of individual mice.

Degeneracy in this system therefore extends to the representational level: the circuit variation produced by individual-target selection is insufficient to leave a detectable footprint in neuronal embedding geometry. Within the framework of Pezon et al. [2026], the circuit structure imposes no detectable constraint on the topology of the similarity-space point cloud (whether measured by Procrustes distance or effective dimensionality). The footprint is absent. To test whether this representational degeneracy is simply a downstream consequence of the structural and functional degeneracy already established, we ran three dissociation tests using permutation Mantel tests on the full 1,431-pair upper triangle of the Procrustes distance matrix. Representational distance was uncorrelated with behavioral distance (Mantel *ρ* = *−*0.007, *p* = 0.871) and with functional sensitivity distance (Mantel *ρ* = *−*0.034, *p* = 0.355), confirming that activity geometry is uncorrelated with the functional axis that individual-target selection does shape. Representational distance showed a weak but significant positive correlation with topological distance (Mantel *ρ* = 0.273, *p <* 0.001; Figure 6B, E, H): circuits sharing more topological scaffold tend toward slightly more similar representations, as expected if structure constrains dynamics to some degree. This residual structure*→*representation coupling does not, however, translate into mouse identity: within-mouse representations are no closer than between-mouse (MW *p* = 0.260), and topology itself carries negligible identity information (NMI = 0.1451; Section 2.4). Representational geometry is therefore degenerate with respect to behavioral identity, even though it is weakly constrained by circuit structure.

Individual-target selection shapes *what* each circuit computes (as shown by the behavioural specialisation and functional sensitivity results) but not *how* that computation is organised in representational space. The geometry of neural activity, like the topology of circuit wiring, is invisible to individual-target selection. Whether this independence extends to the dynamical geometry of the circuit (its stability regime, attractor landscape, and driven trajectory structure) is examined in the following section.

### 2.8 Dynamical geometry is also degenerate

The structural degeneracy established in Sections 2.2–2.6 is most starkly illustrated at the level of individual circuit pairs. Figure 7A shows the positions of two within-mouse pairs within the full topology-behaviour landscape: both lie at near-zero behavioural distance yet are separated by the full topological range of the dataset (pair 1 at the minimum within-mouse Jaccard distance, pair 2 at the maximum). The most topologically similar within-mouse pair in the dataset (mouse D9, replicates 2 and 5; Jaccard distance = 0.694) shares only 11 of 36 edges (30.6%), yet achieves a behavioural distance of 0.0069, effectively producing identical navigation fingerprints. The most topologically dissimilar within-mouse pair (mouse B5, replicates 1 and 5; Jaccard = 0.933) shares only 3 of 45 edges (6.7%), and achieves an equally close behavioural match (0.0068). These two circuits employ architecturally distinct motifs: replicate 1 concentrates connectivity in the recurrent interneuron layer (6 sensory*→*interneuron, 10 interneuron*→*interneuron, 1 interneuron*→*motor connections), while replicate 5 shifts to broader sensory integration with reduced recurrence (11, 4, 3).

**Figure 7.**
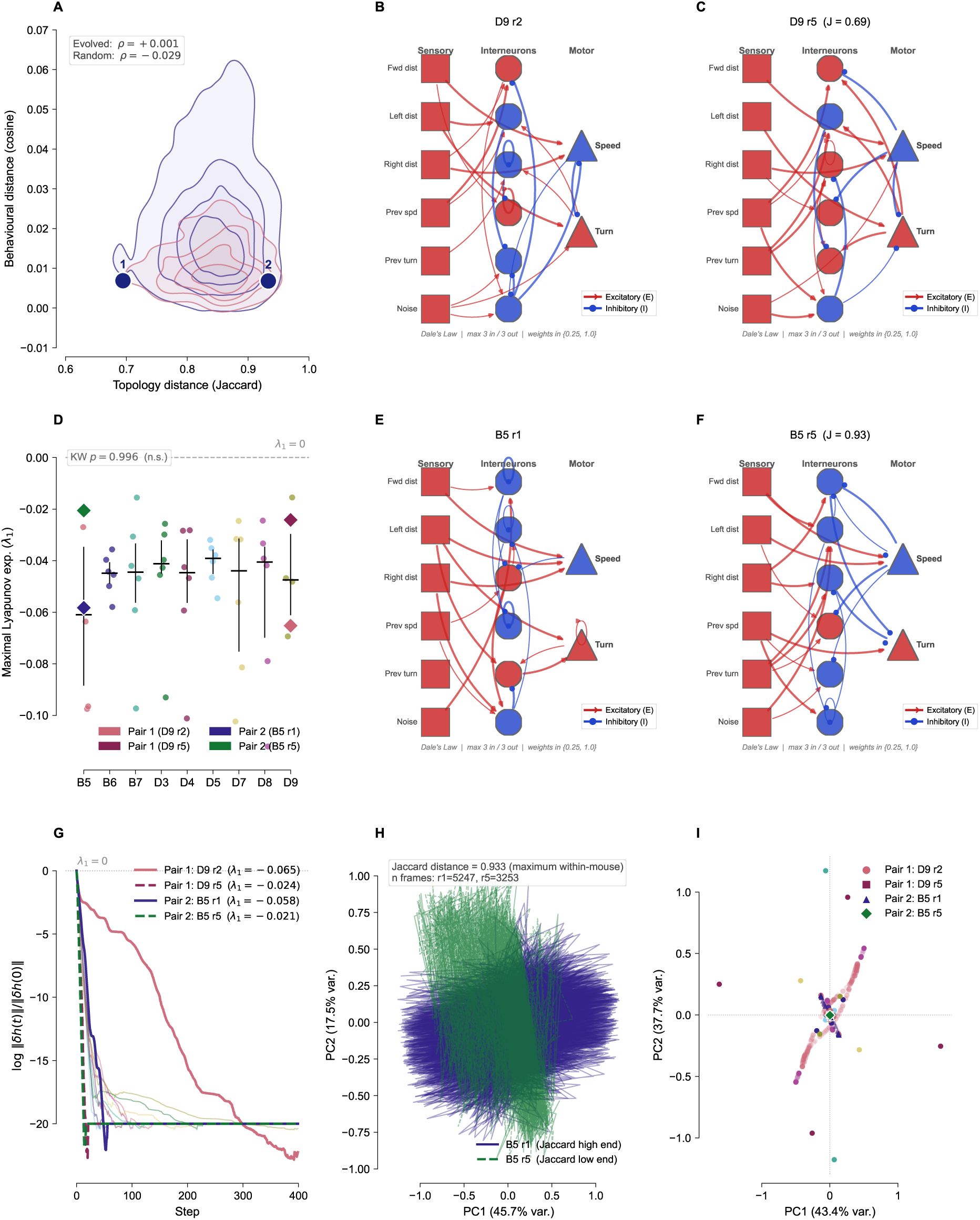
Structural and dynamical degeneracy: topology-disparate circuits are behaviourally and dynamically indistinguishable. Structural and dynamical degeneracy contd…. (A) Joint KDE contours of topology Jaccard distance versus behavioural cosine distance for all 1,431 evolved agent pairs (within-mouse: rose; between-mouse: indigo). Numbered dots mark the two within-mouse pairs selected for circuit comparison: **1** is the most topologically similar pair (mouse D9, Jaccard = 0.694); **2** is the most topologically dissimilar (mouse B5, Jaccard = 0.933). Both lie at near-zero behavioural distance despite spanning the full topological range. **(B, C)** Circuit diagrams for pair 1 replicates (D9 r2 and r5). **(D)** Per-mouse maximal Lyapunov exponent *λ*_1_ for all 54 agents (strip plot with median and IQR); diamond markers indicate the four selected agents; dashed line at *λ*_1_ = 0; Kruskal-Wallis *p* = 0.996 (n.s.). **(E, F)** Circuit diagrams for pair 2 replicates (B5 r1 and r5). Node colours: sensory (squares, blue/red by Dale type), interneuron (circles), motor (triangles). Edge colours: excitatory (red), inhibitory (blue). **(G)** Perturbation decay curves (log ∥*δh*(*t*) ∥ / ∥ *δh*(0) ∥): per-mouse means across all six replicates (thin lines, nine mouse colours) with four selected circuits overlaid in bold; *λ*_1_ values annotated. All circuits collapse toward the same stable regime, with individual trajectories lying within the population envelope. **(H)** Driven trajectory PCA for pair 2 (B5 r1 and r5): interneuron and motor activations (dims 6–13) from all valid maze bouts projected to a shared 2-D PCA. The two circuits – separated by the maximum within-mouse topology distance (Jaccard = 0.933) – trace overlapping trajectories, confirming that trajectory-level degeneracy is topology-independent. **(I)** Autonomous attractor terminal states: all 54 evolved agents run from 200 random initial conditions for 500 steps under zero sensory drive; terminal states projected to a shared 2-D PCA (interneuron and motor dims, colour-coded by mouse). The four selected circuits are highlighted (pair 1: D9 r2 and r5; pair 2: B5 r1 and r5). Population-wide convergence to a common attractor region supports the absence of mouse-specific stability signatures.

**Figure 8.**
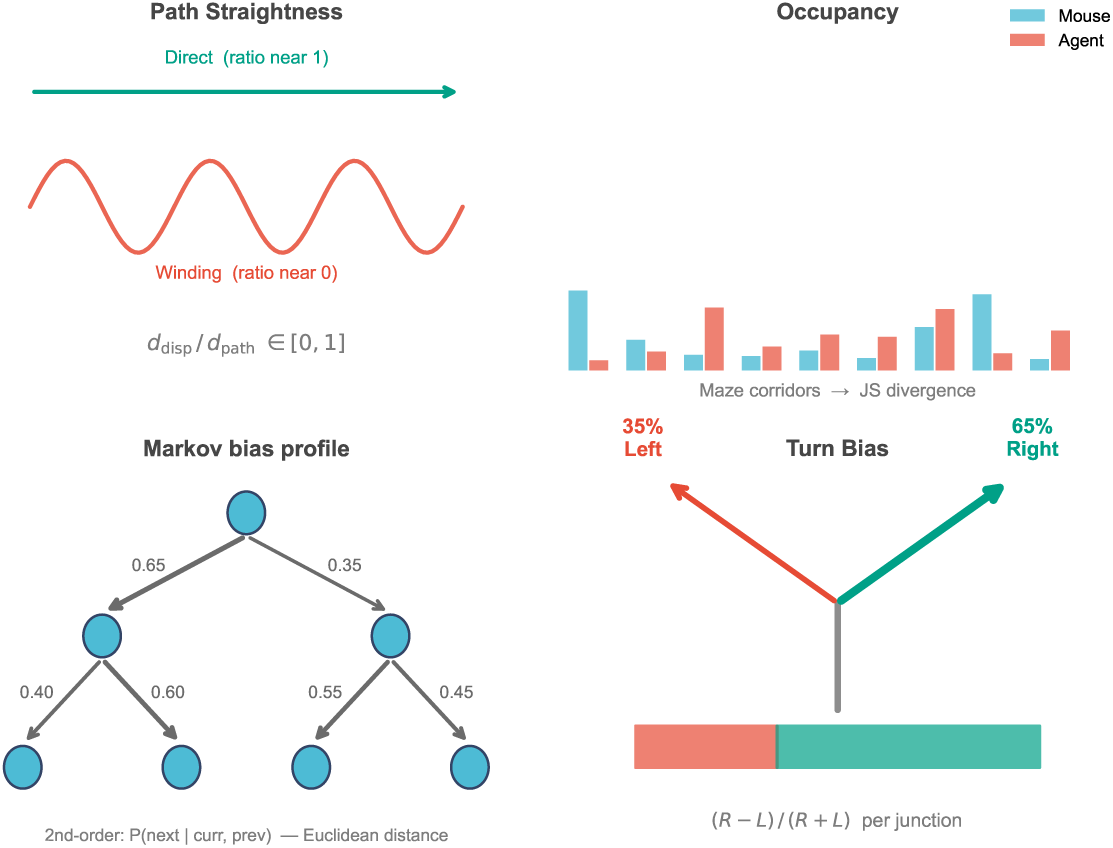
Behavioral fitness metrics. The four components of the fitness function: transition probabilities (Markov bias profile), spatial occupancy (Jensen-Shannon divergence over node-visit distribution), path tortuosity (sliding-window straightness ratio), and turn bias (signed left–right angular velocity asymmetry). Each component is normalized to [0, 1] before weighting.

Extensive structural heterogeneity does not produce corresponding heterogeneity in dynamical character. The maximal Lyapunov exponent *λ*_1_, estimated for all 54 agents via finite perturbation (Section 3.19), was negative for every agent (grand mean *λ*_1_ = *−*0.050 *±* 0.024, range [*−*0.112, *−*0.015]), establishing a uniform slightly-stable dynamical regime across the population. One-way ANOVA across mice returned *F* = 0.345, *p* = 0.943; Kruskal-Wallis confirmed the null (*H* = 1.271, *p* = 0.996): no mouse-specific stability signature exists, despite topological distances spanning the full range from 0.694 to 0.933 (Figure 7D, G).

The degeneracy extends to autonomous dynamics. Each agent was run from 200 random initial conditions under zero sensory drive for 500 steps; virtually all agents converged to a single fixed-point attractor, with no within-versus-between-mouse difference in attractor landscape (sliced Wasserstein distance: within 0.081 *±* 0.125 versus between 0.079 *±* 0.124; MannWhitney *p* = 0.544; Figure 7I).

Driven activity trajectories are likewise degenerate. All 54 agents were presented with an identical 1000-step standardised sensory sequence (Section 3.19); pairwise cosine similarity of mean activation trajectories showed no within-mouse elevation (Mann-Whitney *p* = 0.5085; Supplementary Figure S20B; Figure 7H for the pair-level trajectory PCA).

Across all three dynamical assays, stability regime, attractor landscape, and driven activity geometry, the circuits are indistinguishable. The architectural motif differences visible in Figure 7 (recurrence-heavy versus sensory-integration-heavy wiring) do not produce different dynamical regimes. Degeneracy in this system therefore spans every level examined: aggregate statistics, topology, causal sensitivity, representational geometry, and dynamical geometry.

## 3 Discussion

### 3.1 Degeneracy in constrained circuits: pervasive, structurally invisible, functionally consequential

The central result of this study is a cautionary one for systems neuroscience: behavioral individuation can be real and reliable yet invisible to the entire standard toolkit for relating circuit structure to function. The nine individual targets are genuinely distinguishable—a classifier separates them from the real behavioral data alone at 87% accuracy against a 11% chance level (Supplementary Analysis 3.19), and evolved specialisation persists on held-out trajectories— yet no aggregate statistic, connection-topology measure, representational-similarity or manifold analysis, or attractor/Lyapunov descriptor recovers which mouse a circuit was shaped to become. An analyst equipped with the methods most commonly used to characterise neural circuits would conclude these circuits are interchangeable. They are not.

Our constrained system makes degeneracy an architectural fact rather than an inferred property: the constraint geometry (Dale’s Law, degree limits, quantized weights) compresses all solutions into the same structural region regardless of the behavioral target. This degeneracy has two faces, which we distinguish deliberately. At the level of *competence*, it is classical: the reachable structural space is saturated with equally-good navigators, so selection is free to reach any of them (a many-to-one map from structure to navigational fitness). At the level of *identity*, it is stronger: no structural axis distinguishes which mouse a circuit was shaped to replicate, even though the individual behaviors differ. Individuation exists but leaves no structural trace. We note that not all of the null results carry equal weight: some are close to forced by the architecture. A small saturating tanh network under zero drive contracts to a single stable fixed point, so the absence of a mouse-specific autonomous attractor or Lyapunov signature (§2.8) is largely guaranteed a priori; the non-trivial dynamical test is the driven-trajectory comparison, which is also flat. The structural, topological, and representational nulls are the substantive ones, and it is their conjunction with demonstrable behavioral individuation that constitutes the blind-spot result.

The identity-level degeneracy extends further than the aggregate floor. Connection topology varies freely within and across mice, yet that variation carries no information about behavioral identity at any scale: within mice, between mice, or across the full agent population. The mutual information between any structural axis and behavioral identity is weak and non-significant (maximum NMI = 0.2173). And critically, this topology-behavior flatness is architectural rather than a product of selection: random constrained agents show the same flat relationship as evolved ones (*ρ*_random_ = *−*0.029, *p* = 0.270). Selection does not create the degeneracy; the architecture does. Selection operates within it.

This extends and deepens earlier work on degeneracy in evolved artificial agents. Hu et al. [2025] demonstrated degeneracy at behavioral, structural, and computational levels in Markov brain agents, and showed that it can arise in systems not explicitly designed for it. Our results push this further: in the extreme-constraint regime of 14 neurons, quantized weights, and Dale’s Law, degeneracy operates even on the topology axis itself, the level at which one might most naturally expect individual identity to be encoded. There is no unique “mouse-B5 topology”: the fitness landscape is many-to-one across the entire topological space accessible to the constrained architecture.

Tononi et al. [1999] showed empirically that degeneracy and neural complexity are positively correlated: more degenerate systems tend to be more complex, because degeneracy enables diverse responses to diverse stimuli through a shared structural substrate. Our results are consistent with this picture at the aggregate level but reveal an additional dimension: degeneracy can be present and pervasive even when structural axes are entirely uninformative about behavioral identity. The absence of structural signal does not mean the absence of functional differentiation; it means the differentiation lives somewhere else.

Where it lives is in functional sensitivity commitment. Huang et al. [2024] (preprint) propose a Contravariance Principle: increasing task complexity reduces dynamical degeneracy, because more complex tasks impose tighter constraints on the solution space. Our system inverts this logic by imposing constraint from the architecture rather than the task, and shows that when the architecture saturates the structural axes, selection routes behavioral individuation through a functional channel (sensitivity commitment) that leaves no structural trace. Specialists develop high, varied sensitivity to circuit perturbations, but not convergent: within-mouse sensitivity profiles are no more similar than between-mouse profiles (0.20 versus 0.24; the small gap is not significant under a pseudoreplication-aware test, label-permutation *p* = 0.076), so even replicates of a single mouse do not converge on a shared sensitivity signature. Generalists show substantially lower, more uniformly distributed sensitivity (*σ̄* ^2^ = 0.035) with no strong commitment to any subset of pathways. The 6.4–6.7*×* difference in sensitivity variance between specialists and generalists (mouse-level *p* = 0.002) is the signature of what individual-target selection achieves in a degenerate space: functional commitment without structural differentiation, and without convergence on specific pathways. The representational and dynamical geometry analyses confirm that this absence of convergence is complete: activity embedding geometry (§2.7) and dynamical regime (§2.8) are equally uninformative about mouse identity. Functional sensitivity commitment is the only dimension along which individual-target selection leaves a detectable trace in this system.

### 3.2 The role of architectural constraints

Structural convergence in our system reflects an architectural floor rather than an evolved property: randomly initialised constrained agents return an identical 0/18 significant structural features to evolved agents. The degree limit, quantized weights, and Dale’s Law together reduce the space of realisable circuits so tightly that no aggregate feature can diverge across populations regardless of training. Evolution’s contribution is not convergence of structure (which would happen anyway) but convergence of function: finding computationally specific wiring configurations within that compressed structural space that replicate individual behavioral signatures.

The binary weight scheme *{*0.25, 1.0*}* imposes an analogous compression at the magnitude level. With at most 3 connections per source and only two magnitude values, 61.9% of source neurons carry uniform magnitude assignments that are unaffected by shuffling. Magnitude is therefore a doubly degenerate axis: it carries no information about behavioral identity (NMI = 0.2173) and is causally interchangeable (magnitude permutation reduces MSE rather than elevating it). The architecture forces all variation that matters to live in the one causally potent axis it leaves free: the specific pattern of who-connects-to-whom.

But causal potency is not the same as informational sufficiency. Topology is causally necessary: permuting connection targets disrupts computation (3.53*×* MSE increase). But topological variation is degenerate with respect to behavioral identity: the specific topology an agent uses carries no information about which mouse it was trained on. The architecture makes topology the only axis with causal weight; the fitness landscape makes that axis informationally flat. Selection navigates this constraint by shaping functional sensitivity structure rather than topological pattern. The representational geometry analysis (§2.7) confirms that this structural floor propagates through to the activity level: topology predicts activity embedding geometry (Mantel *ρ* = 0.273, *p <* 0.001), confirming that structure does constrain dynamics, but too weakly and non-specifically to fingerprint behavioral identity at the individual-mouse level.

### 3.3 Implications for neural coding

Our results suggest that aggregate circuit statistics (the metrics most commonly used to characterise neural circuit organisation) may be insufficient to capture functional differences between individuals. Two circuits can have identical aggregate structural profiles (pathway counts, weight magnitudes, E/I composition) while being specialised to distinct behavioral targets. This has direct implications for connectomics efforts that aim to relate circuit structure to function: the mapping from structure to function is many-to-one—many structurally distinct circuits realise the same competence (navigational fitness)—and, more strongly, structure additionally fails to encode *which individual* a circuit implements. We demonstrate this many-to-one, identity-degenerate direction directly; we do not claim the converse plurifunctional (one-structure-to-many-functions) direction, which our design does not test. This matters most in systems where architectural constraints compress the structural space [Bargmann and Marder, 2013, Jonas and Kording, 2017].

The per-source ablation null result (0 of 14 source neurons significant; within-mouse sensitivity profiles no more correlated than between-mouse profiles) extends this to the local level. No single neuron bears disproportionate responsibility for mouse-specific calibration. This holistic distribution parallels ensemble coding in biological systems, where behaviorally relevant information is encoded in distributed patterns of neural activity rather than in the firing of any individual cell [Georgopoulos et al., 1986]. Consistent with this principle, no individual neuron carries disproportionate responsibility for behavioral calibration in our system. Causal weight is distributed across the circuit topology, yet that distributed pattern is itself degenerate across agents.

The functional sensitivity commitment result (A6) suggests a specific prediction for biological circuits. If the principle generalises, one would expect circuits evolved or trained for individual-specific behavioral targets to show higher, more consistent sensitivity to perturbations of particular pathways than circuits optimised for population-level or multi-target performance, even when the circuits are structurally indistinguishable by any aggregate measure. The representational degeneracy result (§2.7) deepens this prediction: functional individuation leaves no trace even in the geometry of neural activity embeddings, suggesting that population-activity analyses commonly used in systems neuroscience (manifold dimensionality, cross-session RSA, CKA) may be systematically insensitive to individual-specific circuit organisation even when that organisation is strong. Measuring functional sensitivity commitment (rather than structural or representational similarity) may therefore be a more productive strategy for identifying individual-specific circuit organisation in biological neural systems.

### 3.4 Limitations

Several limitations apply. First, the quantized weight scheme (*{*0.25, 1.0*}*) limits the resolution of synaptic strength tuning. Continuous weights might reveal more pathway-level magnitude variation and could alter the topology-vs-magnitude result, which may partly reflect the architecture’s unusually compressed magnitude space rather than a general principle [Huang et al., 2024].

Second, the fitness function captures behavioral statistics rather than moment-by-moment trajectory matching, and the behavioral fingerprint (4 metrics from maze navigation) may not represent all relevant dimensions of individual mouse identity. Other behavioral contexts, additional metrics, or richer trajectory representations might reveal structural signals invisible to the current fitness function.

Third, the 14-neuron network is a deliberate minimal model. Real navigation circuits in rodents involve millions of neurons across hippocampus, entorhinal cortex, and striatum. Whether topology-dominant causal encoding and functional sensitivity commitment as the individuating property scale to larger, biologically realistic networks is an open question. Degeneracy and neural complexity are empirically positively correlated [Tononi et al., 1999], suggesting structural convergence may become more pronounced in larger networks; the topology-behavior flatness we observe, however, emerges already in the minimal 14-neuron case and is architectural rather than size-dependent.

Fourth, the sensitivity RSA analysis (A4) is likely underpowered in the current dataset. The Mantel test and within/between MW statistic are both null (Mantel *ρ* = *−*0.044, permutation *p* = 0.357), but with 54 agents across 9 mice the geometric resolution of the 14-dimensional sensitivity space is limited. Whether the *geometry* of the sensitivity profile (as opposed to its variance) carries individual-specific information at higher replicate counts remains an open question. Additional replicates (targeting *n* = 10 per mouse) are planned. More generally, with six replicates per mouse a non-significant between-mouse contrast could in principle reflect limited statistical power rather than genuine equivalence. The structural, representational, and dynamical degeneracy nulls are guarded against this because each is paired with a matched random-agent architectural control that reproduces the same null—locating the uniformity in the constrained architecture rather than in insufficient sampling—while the aggregate-feature (0/18) null is additionally backed by the large-effect power analysis (Supplementary Figure S7).

Fifth, results are drawn from 9 mice across two inbred strains navigating a single maze geometry without reward. Generalisation to outbred populations, different maze conditions, or different task structures requires further investigation. These findings constitute a proof-of-concept demonstration in a controlled minimal system, not a general claim about neural coding mechanisms.

Sixth, the analysis is null-heavy by design, and the interpretation rests on a small number of positive results, each tied to a specific pre-specified hypothesis rather than emerging from an unstructured screen: the topology-to-representation correspondence (Mantel *ρ* = 0.273, *p* =*<* 0.001) and the specificity dose-response (Spearman *ρ* = 0.68, permutation *p* = 0.050, *n* = 9). We report these with effect sizes and confidence intervals rather than *p*-values alone, and do not apply a joint multiple-comparison correction across them because they test distinct, independently motivated hypotheses. The dose-response in particular is borderline and, as noted above, indicative rather than firmly established; neither the causal nor the representational conclusion depends on it.

### 3.5 Future directions

Several extensions would strengthen and generalise these findings. First, increasing replicates to 10–15 per mouse would provide adequate power for the sensitivity RSA geometry analysis (A4) and would also improve resolution on medium-effect structural features. This is tractable within the current evolutionary framework.

Second, relaxing architectural constraints, i.e. allowing more connections per neuron, continuous weights, or removing Dale’s Law, would test whether the topology-behavior flatness and functional sensitivity commitment results depend on the extreme-constraint regime or persist across a wider range of architectures. The representational (§2.7) and dynamical (§2.8) geometry analyses provide additional comparison axes: whether representational and dynamical degeneracy persist in less constrained circuits, or whether structural signal propagates through to activity geometry and stability regime at higher architectural complexity.

Third, the generalist result (Section 2.6) opens a productive experimental direction: training agents on subsets of mice (2, 3, or 4 targets) would allow systematic characterisation of how functional sensitivity commitment degrades as a function of the number of behavioral targets, and at what point individual-target specificity becomes impossible to maintain.

Fourth, applying the same framework to other behavioral datasets, i.e. different species, tasks, or maze geometries, would test whether functional sensitivity commitment as the individuating property is specific to mouse navigation or a more general consequence of individual-target optimisation in constrained circuit spaces.

Finally, bridging to real neural recordings would test the biological relevance of the mechanisms identified here. The computational circuit predicts that individually-calibrated circuits should show higher functional sensitivity commitment than population-optimised ones, even when structurally indistinguishable. This prediction is testable using perturbation experiments (optogenetic or pharmacological silencing of identified pathway types) combined with behavioral readout in animals with known individual behavioral profiles.

## Methods

### 3.6 Mouse behavioral data

We used behavioral data from 9 mice (B5, B6, B7, D3, D4, D5, D7, D8, D9) exploring a hierarchical binary tree maze without reward, from Rosenberg et al. [2021]. For each mouse, we computed four behavioral metrics from the raw trajectory data:

1. Transition probabilities between maze arms (Markov property): Decision-making at various nodes in the binary tree representing the maze can be characterised with a Markov-chain analysis. For each node in the maze, we compute the probability distribution over next-node transitions based on the direction of entry (from the bottom, left or right arm), and that of left versus right, yielding a 6-dimensional vector per node. These are averaged across a level and stacked across the tree, both for the agent and the mouse. The Euclidean distance between these bias profiles becomes the penalty for the agent.
2. Spatial occupancy as a node-visit distribution: A probability distribution over the maze’s corridor nodes (runs) is constructed by weighting each node proportionally to the total number of frames the mouse spent there within a bout. Dwell time for each node is computed as the difference between consecutive start-frame entries in the Rosenberg bout traces, matching the reference implementation of Rosenberg et al. [2021]. The Jensen-Shannon divergence between the agent and mouse node-occupancy distributions is applied as a penalty, encouraging the agent to explore the maze with similar spatial coverage as the mouse.
3. Path tortuosity: Path efficiency is measured using sliding window straightness. For each window, we compute the ratio of the displacement to distance. A penalty is applied according to relative deviation from the mouse baseline.
4. Turn bias: A simple turn bias (left versus right) is calculated using the distribution of angular velocities. The turn bias is the signed left–right asymmetry (n_right *−* n_left)/n_total *∈* [*−*1, 1], and the agent is penalised for deviation from the mouse’s turn bias. Simulated evaluation consists of 20 independent bouts of 2000 frames each.

### 3.7 Network architecture

Each agent is a recurrent neural network with 14 neurons: 6 sensory (front, left, right wall distances; previous speed; previous turn rate; Gaussian noise), 6 interneurons, and 2 motor (speed, turn rate). The network is updated at each timestep according to:

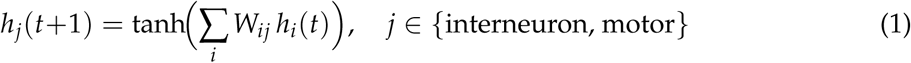

where **W** is the 14 *×* 14 weight matrix with *W_ij_*denoting the connection from neuron *i* to neuron *j*, and **h** is the state vector. In matrix notation, **h**(*t* +1) = tanh(**W***^⊤^* **h**(*t*)), equivalently tanh(**h**(*t*)*^⊤^***W**) for **h** a row vector—the form used in the reference implementation (state @ W), with *W_ij_* the weight from *i* to *j* throughout. Sensory neurons are clamped to their input values at each step; only interneuron and motor states are updated dynamically.

Three biological constraints are enforced:

1. **Dale’s Law**: Each neuron is assigned a fixed type (excitatory or inhibitory). All outgoing weights share the source neuron’s sign: *w_ij_* = *|w_ij_| ·* type*_i_*. Sensory neurons are always excitatory (consistent with the predominantly glutamatergic nature of sensory afferents [Kandel et al., 2021]); interneuron and motor types are evolved.
2. **Sparse connectivity**: Each neuron has a maximum of 3 incoming and 3 outgoing connections. Sensory neurons receive no incoming connections. All other connection types are permitted: sensory*→*interneuron, sensory*→*motor, interneuron*→*interneuron, interneuron*→*motor, and motor*→*interneuron (recurrent feedback). Self-connections (autapses) are permitted; empirically, 80% of evolved agents (43 of 54) acquired at least one self-connection on interneurons or motor neurons.
3. **Quantized weights**: Magnitudes are drawn from *{*0.25, 1.0*}*. Combined with Dale’s Law, the effective weights take values in *{−*1.0, *−*0.25, 0, +0.25, +1.0*}*.

### 3.8 Simulation environment

The agent navigates a hierarchical binary tree maze rendered as a 2D polygon. At each timestep, sensory inputs include 3 raycasts (front, left, right wall distances), previous speed, previous turn rate, and noise (*N* (0, 0.1)). Motor outputs are filtered through momentum dynamics:

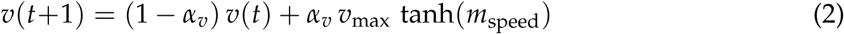

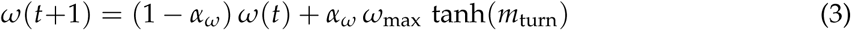

where *α_v_* = 0.263 and *α_ω_* = 0.437 are momentum coefficients, and *v*_max_ = 1.949 and *ω*_max_ = 1.191 are the maximum speed and turn rate, all derived empirically from cross-subject medians of mouse trajectory statistics (lag-1 autocorrelations for speed and turn rate; 99th-percentile speed and turn distributions; Gaussian smoothing *σ* = 1.0 applied prior to parameter estimation). Full derivation details are provided in Supplementary Methods. Each evaluation consists of 20 bouts of 2000 frames.

### 3.9 Evolutionary algorithm

We use a (*µ* + *λ*) evolutionary strategy with a population size of 500 and an elite fraction of 10% (50 agents). Three mutation operators are applied to offspring:

1. Weight magnitude toggle (probability *p_w_*): flips a connection’s magnitude between 0.25 and 1.0
2. Structural rewiring (probability *p_s_*): adds, removes, or redirects a connection
3. E/I type flip (probability *p_n_*): toggles a neuron between excitatory and inhibitory

Mutation rates follow a linear decay from 15% to 1% over 200 total generations. All results are reported at generation 150, where population fitness had plateaued in all 54 runs (defined as *<*2% change across the final 30 generations); at this checkpoint the mutation rate was approximately 4.5%, which did not affect outcomes as fitness had plateaued by generation 100–120 for all runs. The remaining 50 generations produced negligible improvement. A connectivity repair step re-enforces structural constraints after each mutation: (i) per-neuron degree limits (max 3 in/out) by pruning excess connections uniformly at random; (ii) no incoming connections to sensory neurons; (iii) minimum 1 outgoing connection per non-motor neuron. Self-connections (autapses) are not prohibited and frequently arise during evolution (see Section 3.7).

### 3.10 Fitness function

Fitness is defined as:

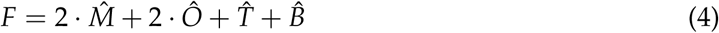

where each component is normalized to [0, 1] using fixed reference constants derived from the behavioral range of random agents: *M*^^^ = min(1, *d*^Markov^_Eucl_ /6), *O*^^^ = min(1, *d*^Occupancy^_JS_), *T*^^^ = min(1, *|*Δtortuosity*|*/tortuosity_mouse_), and *B*^^^ = min(1, *|*Δturn bias*|*). The Occupancy component uses Jensen-Shannon divergence; the Markov component uses Euclidean distance between agent and target bias profiles. Tortuosity and Turn bias use differences from target scalar values. Lower fitness indicates a better match to the target mouse. The 2*×* weighting on Markov and Occupancy reflects that these distributional measures (JS divergence over decision-making transition patterns and spatial coverage) are richer in individual-specific information than the scalar statistics captured by Tortuosity and Turn bias. The sensitivity of this fitness function to individual mouse differences is validated by the cross-mouse generalization result (Section 2.2): agents optimized for one mouse achieve substantially higher fitness errors on other mice’s behavioral profiles (off-diagonal mean 1.122 *±* 0.242 vs diagonal 0.748 *±* 0.113).

### 3.11 Circuit feature extraction

For each of 54 best-evolved agents, we extracted 18 structural features spanning four categories: connectivity (total connections, density), E/I composition (ratio, excitatory/inhibitory counts), pathway counts (SI, SM, II, IM, MI, MM), E/I fractions per pathway, interneuron degree statistics, and weight statistics (mean magnitude, fraction at maximum). A baseline of 200 random agents was generated for comparison.

### 3.12 Statistical analysis

#### Per-mouse ANOVA

One-way ANOVA with Benjamini-Hochberg FDR correction (*α* = 0.05) tested 18 features across 9 mice (*k* = 9, *n* = 6 per group). Permutation tests (10,000 permutations) provided non-parametric validation. BH-FDR was preferred over familywise-error correction (e.g., Bonferroni) because it controls the expected proportion of false discoveries rather than the probability of any false discovery, providing greater power for this discovery analysis across 18 features. The FDR guarantee holds under positive dependence among test statistics [Benjamini and Hochberg, 1995], which applies here as pathway counts co-vary with total connection count. The null result is robust to correction method: although the smallest raw *p*-value is 0.045 (interneuron fan-in), it does not survive correction (*p*_FDR_ = 0.476), the permutation test agrees (*p*_perm_ = 0.051), and no feature is significant after BH-FDR (0/18, all *p*_FDR_ *>* 0.47).

#### Evolved vs random comparison

One-sample *t*-tests with Cohen’s *d* and bootstrap 95% CIs (10,000 resamples).

#### Power analysis

Post-hoc power for the Wilcoxon signed-rank permutation specificity test (*n* = 9 paired observations, one-tailed *α* = 0.05) was computed via the normal approximation to the Wilcoxon distribution. The observed effect size was Cohen’s *d_z_* = 1.755, yielding attained power *>* 0.999. The minimum detectable effect at 80% power was *d_z_* = 0.829 (0.209 fitness units); the observed effect exceeds this threshold by a factor of 2.12, making the *n* = 9 design adequate (Figure S8). For the circuit-level and structural feature analyses, achieved power was computed via the non-central *F* distribution and minimum detectable *η*^2^ was determined by binary search.

#### Cross-mouse generalization

A 9 *×* 9 fitness matrix was computed by evaluating each mouse’s best agent on all 9 behavioral targets. The behavioral specialization index is defined as 1*−* (diagonal mean / off-diagonal mean); a value of 0 indicates no specialization, positive values indicate specialization. The raw own/cross ratio equals 1*−* index.

#### Strain clustering

Mann-Whitney *U* test comparing within-strain vs between-strain off-diagonal fitness values.

#### Weight similarity

Pairwise cosine similarity of flattened weight matrices. Within-mouse vs between-mouse comparison via Mann-Whitney *U*.

#### Activation trajectory analysis

All 54 best agents were run on an identical 1000-step standardized sensory input sequence (random seed fixed at 42). Only the three wall-distance inputs (*d _f_* , *d_l_*, *d_r_*) follow a sinusoidal profile; the speed and turn proprioceptive inputs are derived from the agent’s own motor outputs at each step (closed-loop feedback), consistent with how agents operate during actual maze navigation. This sequence does not come from actual maze trajectories; it is a synthetic standardized probe designed to elicit comparable dynamics across all agents. Hidden-unit states (8 dimensions: 6 interneurons + 2 motors) were recorded at each timestep.

Two representational similarity matrices (RSMs) were computed: (1) pairwise cosine similarity of per-agent mean activation vectors, and (2) mean pairwise Pearson correlation across all 8 dimensions of the full 1000-step trajectory. Within-vs between-mouse similarity was compared via Mann-Whitney *U* (one-sided, *α* = 0.05). PCA was applied to mean activation vectors for visualization.

Maximal Lyapunov exponents were estimated for all 54 agents via finite perturbation (*ε* = 10*^−^*^6^, 8 random unit-vector perturbation directions). Log divergence Λ(*t*) = log(∥ **h**(*t*) *−* **h***_ε_*(*t*) ∥ /*ε*) was tracked for 400 steps; *λ*_1_ was estimated as the mean log-divergence rate over the full 400-step trajectory (Equation 7), averaged over the 8 perturbation directions. One-way ANOVA and Kruskal-Wallis tests compared *λ*_1_ across mice.

#### Fixed point analysis

For each agent’s 8*×*8 autonomous submatrix **W**_sub_ = **W**_[6:14,6:14]_ (interneurons and motors under zero sensory drive, i.e. the circuit’s intrinsic attractor structure independent of input), fixed points were found via iterative convergence (*h ←* tanh(*h* ← tanh(**W**^⊤^_sub_*h*), 300 random restarts, tolerance 10*^−^*^5^, maximum 5000 iterations), consistent with the forward-pass convention *h_j_* = tanh(∑*_i_ W_ij_h_i_*). Duplicate fixed points were removed at distance *<* 10*^−^*^3^. The Jacobian at each fixed point was computed as **J** = diag(1 *−* (**h***^∗^*)^2^) *·* **W**^⊤^_sub_, and the dominant eigenvalue *|λ*_max_*|* was obtained via scipy.linalg.eigvals. One-way ANOVA and Kruskal-Wallis tests compared fixed point count and *|λ*_max_*|* across mice.

#### Weight permutation ablation

For each of 54 agents, 20 permuted variants were generated using a source-preserving scheme. Within each of 6 pathway blocks (SI, SM, II, IM, MI, MM), we permuted only which target neurons each source connects to, keeping per-source outdegree, weight magnitudes, and Dale’s Law sign fixed. By *connection topology* we mean precisely this assignment of source-to-target pairs within each pathway block: which *k* of the available target neurons each source connects to, given fixed out-degree, Dale’s Law sign, and weight magnitudes. This is distinct from graph-theoretic topology (cycle structure, path lengths, spectral properties), which is preserved by construction since the degree sequence and block structure are unchanged. This preserves all 18 structural features exactly, including E/I fractions per pathway block.

Each evolved agent’s permutation space is large: the mean number of distinct connectivity assignments consistent with the structural constraints is 10^12.5^ configurations (range 10^11.2^– 10^14.0^ across 54 agents; Supplementary Analysis). Of source-pathway combinations, 59% have non-trivial permutation freedom (*>*1 available target assignment). Sensory-to-interneuron connections carry the most freedom (SI: mean *C* = 9.5 *±* 2.2 choices per source); motor-target pathway blocks are near-deterministic (SM: *C* = 1.5, IM: *C* = 1.3, MM: *C* = 1.3), reflecting the extreme sparsity of motor connectivity. Preservation was verified programmatically: Max *|*Δfeature*| <* 0.001 across all 18 features for all tested agents.

Motor output MSE between original and permuted agents was computed on the same standardized 1000-step input sequence. Mann-Whitney *U* (one-sided) compared the permutation MSE distribution to same-mouse replicate-pair MSE, which serves as the natural within-mouse variability floor.

### 3.13 Mouse-specific permutation fitness test

For each of 9 mice, the best-evolved agent (minimum fitness across all 6 replicates at generation 150) was selected. Ten source-preserving permuted variants were generated per agent (rationale below). The original and all 10 permuted agents were evaluated against each of the 9 mice’s behavioral baselines using the full simulation pipeline (20 bouts *×* 2000 frames per evaluation).

Fitness degradation was computed as Δfit = fitness_permuted_ *−* fitness_original_ for each (agent, permutation, test-mouse) triple. Δfit_own_ is the mean degradation when the test mouse matches the training mouse; Δfit_other_ is the mean over the 8 non-training mice.

This number of permutations was chosen to balance computational cost (each evaluation requires 20 bouts *×* 2000 frames *×* 9 behavioral targets per agent, totalling *∼*360,000 simulation steps per permutation) against statistical resolution. The pooled Mann-Whitney test aggregates *n* = 90 own-mouse and *n* = 720 other-mouse Δfit observations across all 9 mice, providing adequate power. With only *n* = 10 paired observations per mouse, the within-mouse Wilcoxon signed-rank test is near its minimum achievable *p*-value (*∼* 0.001); the pooled Mann-Whitney *U* is the primary statistic.

Statistical tests: pooled Mann-Whitney *U* (one-sided: Δfit_own_ *>* Δfit_other_); Wilcoxon signed-rank on the 9 per-mouse paired means (Δfit_own_ *−* Δfit_other_, *n* = 9, one-tailed) as confirmation. The D5 analysis used D5’s single best agent (*n* = 10 permutations); no within-agent significance test is possible with *n* = 1 agent.

### 3.14 Magnitude-only permutation

To isolate the contribution of connection topology vs synaptic strength, we implemented a magnitude-only permutation. For each source neuron, the connection targets (topology) are held fixed, and only the *{*0.25, 1.0*}* magnitude assignments are shuffled among that source’s existing connections. Dale’s Law sign is preserved. Motor output MSE was computed as for the topology permutation (*N* = 20 permutations per agent, same 1000-step sensory sequence). The replicate baseline (1.0*×* by definition) serves as the null expectation.

### 3.15 Per-source ablation sensitivity analysis

For each of 54 best-evolved agents and each of 14 neurons (indices 0–13), a source-ablated agent was generated by permuting only that neuron’s outgoing connections within its relevant pathway blocks, keeping all other connections unchanged. Motor output MSE against the original agent was computed on a fixed 500-step sensory sequence (random seed 99). Raw MSE values were normalized by each agent’s full-circuit permutation baseline MSE, yielding a 54 *×* 14 relative sensitivity matrix.

Statistical analysis: one-way ANOVA per source neuron across 9 mice (sensitivity as dependent variable, mouse identity as factor; Bonferroni correction for 14 simultaneous tests). Sensitivity vector correlation: Pearson *r* between all pairs of agents’ 14-dimensional sensitivity vectors, classified as within-mouse (same training mouse, different replicates) or between-mouse. Mann-Whitney *U* compared the two distributions (one-sided, within *>* between).

### 3.16 Topology-behavior correlation analyses (A1–A5)

To quantify the relationship between structural variation and behavioral identity, we computed pairwise Spearman correlations between topology distance and behavioral distance across all agent pairs. Topology distance was measured as Jaccard distance between binarised weight matrices (connection presence/absence). Behavioral distance was measured as cosine distance between each agent’s row of the 54 *×* 9 fitness matrix. Analysis A1 computed these correlations across all (^54^) = 1431 agent pairs and separately within and between mouse groups. Analysis A2 restricted to within-mouse replicate pairs (*n* = 135) and additionally computed partial Spearman correlations controlling for magnitude and sign distance. Analysis A3 quantified mutual information (MI) and normalised mutual information (NMI) between each structural axis and mouse identity using *k*-means clustering (*k* = 5) as a discretisation step, following the framework of Tononi et al. [1999]. We chose *k* = 5 as a number of clusters well-supported by 54 agents (*≈*11 per cluster on average), providing resolution to detect structural grouping without overfitting noise. To confirm the null result is not an artefact of this choice, we additionally computed adjusted mutual information (AMI), which corrects for chance-level agreement and is independent of cluster count. AMI was near zero or negative for all three axes across *k* = 3– 9 (topology: *−*0.06–0.05; magnitude: *−*0.03–0.04; sign: *−*0.06–0.02), confirming that the null holds regardless of discretisation choice. Analysis A4 performed representational similarity analysis (RSA) on the 54 *×* 14 per-source ablation sensitivity matrix, computing Mantel test correlation between the resulting 54 *×* 54 sensitivity RSM and a mouse-identity RSM (within = 1, between = 0); permutation *p*-values used 10,000 random permutations. Analysis A5 repeated the topology-behavior correlation on 54 randomly initialised (unevolved) constrained agents to establish the architectural null.

### 3.17 Generalist vs specialist sensitivity comparison (A6)

To compare functional sensitivity commitment between individual-target specialists and generalist agents, we computed per-neuron sensitivity variance across replicates. For each of the 14 source neurons, we recorded the ablation sensitivity (MSE increase under individual source permutation) for all replicates of each agent type. Raw MSE values were first normalised by each agent’s full-circuit permutation baseline MSE (the expected MSE under random topology permutation of all connections simultaneously, estimated from the *N* = 20 full permutations used in the topology permutation analysis), yielding a relative sensitivity score that is comparable across agents regardless of each agent’s baseline computational output scale. Sensitivity variance per neuron was defined as the variance of these normalised sensitivity scores across the replicate set. For specialists, this variance was computed across the 6 replicates per mouse, then pooled across all 9 mice; for generalists, across the 15 generalist replicates. The within-mouse-pooled specialist:generalist ratio is 6.4*×*; because between-mouse variance is negligible, computing the specialist variance across all 54 agents instead gives 6.7*×*, so we report the range 6.4–6.7*×*. Because per-neuron variances within a circuit are not independent, the primary test uses the mouse as the unit (one-sample Wilcoxon on the 9 per-mouse commitment scalars against the generalist: 9/9 mice exceed it, *p* = 0.002); a hierarchical bootstrap (resampling mice, replicates, and generalists; seed 1) gives a 95% CI of [2.88, 11.92]. The per-neuron Mann-Whitney (*n* = 14 per group, one-sided specialist *>* generalist; *p* = 0.001) is reported for comparability but is pseudoreplicated. Topology cosine similarity was compared between groups using a two-sided Mann-Whitney *U* test. Generalist fitness cost per mouse was computed as the percentage difference between mean generalist fitness and specialist fitness on the same mouse’s behavioral target.

### 3.18 Representational geometry analysis (B1–B5)

To test whether individual-target selection produces convergence in neuronal embedding space, we re-simulated all 54 best-evolved agents under a fixed random seed (seed = 42) for 20 independent maze bouts of up to 2,000 steps each (identical maze geometry and physics parameters as fitness evaluation). The full 14-dimensional network state was recorded at every timestep. For each agent, all valid activity frames across bouts were concatenated to form a 14 *× T*_total_ activity matrix, where *T*_total_ is the sum of exit-frame counts across bouts. Each neuron’s time series was independently standardised to zero mean and unit variance.

#### PCA loadings

PCA (6 components) was applied per agent to the transposed activity matrix (*T*_total_ *×* 14). The *loading matrix* **L** *∈* R^14^*^×^*^6^ (columns = PC directions in neuron space, i.e. pca.components_.T) constitutes the neuronal embedding [Pezon et al., 2026]: neurons whose activity covaries across maze bouts cluster in loading space. Because each PC direction has unit norm by construction, the columns of **L** are orthonormal. The sign ambiguity of independent PCA solutions was removed by aligning each agent’s loading matrix to the first agent: any PC column whose dot product with the corresponding reference column was negative was sign-flipped.

#### Manifold dimensionality

The effective dimensionality at the 90% variance threshold was defined as the smallest *k* such that ∑*^k^*_*i*_=1 λ_*i*_ ≥ 0.90, where λ_1_ ≥ … ≥ λ_6_ variance ratios. The participation ratio

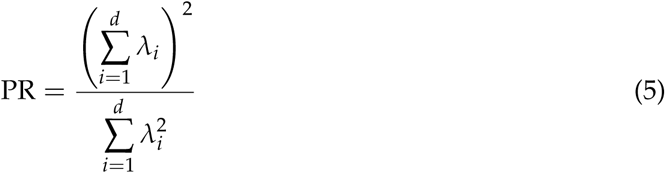

provides a continuous effective dimensionality ranging from 1 (one dominant component) to *d* = 6 (all components contribute equally).

#### Procrustes distance

Pairwise embedding similarity was quantified via orthogonal Procrustes alignment. For agents *a* and *b* with loading matrices **L***_a_*, **L***_b_ ∈* R^14^*^×^*^6^ (orthonormal columns), we found the rotation **R***^∗^* minimising ∥**L***_a_ −* **L***_b_***R∥** *_F_* using scipy.linalg.orthogonal_procrustes, which returns the alignment quality *s* = tr(**L**^⊤^_*a*_**L**_b_**R**^*^) (sum of singular values of **L**^⊤^_*a*_**L**_b_). The Procrustes distance equals the residual Frobenius norm after optimal alignment:

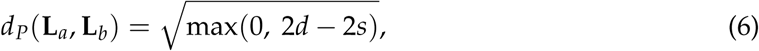

where *d* = 6 is the number of retained components. This distance is zero when the two loading matrices are identical up to rotation, and reaches *^√^*2*d* when they span orthogonal subspaces.

#### Mantel tests

To test whether representational distance correlated with behavioral, topological, or functional sensitivity distance, we applied permutation Mantel tests (9,999 permutations) to the upper triangle (*n* = 1,431 agent pairs) of the 54 *×* 54 Procrustes distance matrix. The test statistic was Spearman *ρ* between the two distance vectors; the reported *p*-value is the fraction of permuted *ρ* values exceeding the observed value.

### 3.19 Dynamical geometry analysis (§2.8)

#### Maximal Lyapunov exponent

For each of the 54 best-evolved agents, the maximal Lyapunov exponent *λ*_1_ was estimated via finite perturbation. The agent was initialised under the standardised synthetic sensory sequence used throughout (identical to the sequence in Section 3.13), and *N*_dir_ = 8 random perturbation directions of magnitude *ε* = 10*^−^*^6^ were applied to the initial network state. Each perturbed trajectory was evolved for 400 steps under the same sensory input, and the log-ratio of perturbation norms at each step was recorded. The Lyapunov exponent was obtained as the mean log-divergence rate over the full 400-step trajectory, averaged over the 8 perturbation directions:

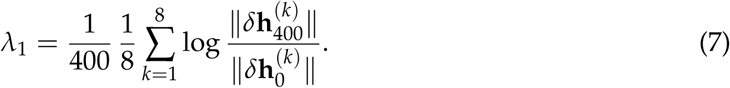

Between-mouse differences in *λ*_1_ were assessed with one-way ANOVA (9 groups) and confirmed with Kruskal-Wallis.

#### Attractor landscape

Each agent was initialised from 200 random network states drawn uniformly in [*−*1, 1]^14^, with zero sensory input, and evolved for 500 steps. The distribution of terminal states was used to characterise the attractor landscape. Pairwise landscape dissimilarity was quantified as sliced Wasserstein distance (50 random projections) between the two sets of terminal states. The 54 *×* 54 distance matrix was partitioned into within-mouse and between-mouse pairs; a Mann-Whitney *U* test compared the two distributions.

#### Trajectory RSM

All 54 agents were run on the same 1,000-step standardised sensory input sequence (Section 3.13). The mean activation vector was computed for each agent by averaging the 14-dimensional network state across all 1,000 steps, yielding a single population-mean representation per agent. Pairwise cosine similarity between all 1,431 agent pairs formed the trajectory representational similarity matrix (RSM). Within-mouse and between-mouse similarity distributions were compared with a Mann-Whitney *U* test.

## Data availability

All code, the distilled best-evolved agents, the analysis intermediates, and the scripts that regenerate every figure and statistic reported here are available at https://github.com/pranetkhetan/ constrained-neuroevolution and are archived on Zenodo (DOI: 10.5281/zenodo.21949631). These include the aggregate individuation result (analysis/individuation_results.pkl) that underlies the classifier accuracy reported in the Discussion and Supplementary Analysis 3.19. This code and analysis archive alone is sufficient to reproduce every result reported here; no large download is required. The same Zenodo record additionally contains the full evolutionary archive that these artifacts were distilled from—all 150 generations of the 54 specialist and 15 generalist runs, together with the full per-frame activation tensors—which is needed only to re-derive the analysis intermediates from scratch or to audit the evolutionary trajectories directly. Raw mouse behavioral data are the tracked trajectory (-tf) files from Rosenberg et al. [2021] (eLife 2021; doi:10.7554/eLife.66175), which we do not redistribute. To regenerate the individuation result from source, obtain those files, place them as data/raw/{mouse}-tf for the nine mice, and run python scripts/analyze_individuation.py --tf_dir data/raw--n_folds 6 --fold_mode stride (deterministic, seed = 1); all other results regenerate from the shipped intermediates via python scripts/build_paper_stats.py.

## Acknowledgments

We thank the Ashoka University Lodha Genius Program, 2025, for support. Computation was carried out on Google Colab and personal workstations. We are grateful to the authors of Rosenberg et al. [2021] for making their data publicly available.

## Author Contributions

PK: Conceptualization, Investigation, Writing – Original Draft. AA: Conceptualization, Formal analysis, Investigation, Visualization, Writing – Review & Editing. Both authors approved the final version.

