## Supplementary Material for "Selection navigates a degenerate circuit space: behavioral individuation without structural differentiation in constrained neuroevolution"

Pranet Khetan
Shiv Nadar School, India

Aditya Asopa, PhD
Visiting Faculty, Lodha Genius Program 2025, Ashoka University, India
Research Scientist, Anthriq, India

**Abstract**

This document contains the Supplementary Information for the manuscript "Selection navigates a degenerate circuit space: behavioral individuation without structural differentiation in constrained neuroevolution." The study evolves 14-neuron recurrent circuits under three biological constraints (Dale's Law, sparse connectivity, and quantized weights) to replicate the natural navigation behavior of nine individual mice, and shows that behavioral individuation is robust yet leaves no legible trace on any aggregate structural, topological, representational, or dynamical axis; the single axis that individual-target selection shapes is the *strength* of functional sensitivity commitment. This Supplementary Information provides the supporting control analyses and figures referenced throughout the main text — convergence and network structure, emergent behavior and excitatory/inhibitory dynamics, statistical power and specialization controls, structural- and sensitivity-null analyses, and representational- and dynamical-geometry robustness — comprising Supplementary Figures S1–S23 and Supplementary Tables S1–S2.

**Reference to main text.** Section, figure, and table numbers without an "S" prefix (e.g. §2.6, Figure 5) refer to the main manuscript; all items with an "S" prefix are contained in this document.

### Supplementary Analysis

#### S1. Per-mouse individual replicate trajectories

Population mean and best-of-generation fitness for each of the 9 mice across 150 generations. Each panel shows a single mouse; 6 individual replicate trajectories are shown. All runs converge by generation 100–120 with no aberrant behaviour.

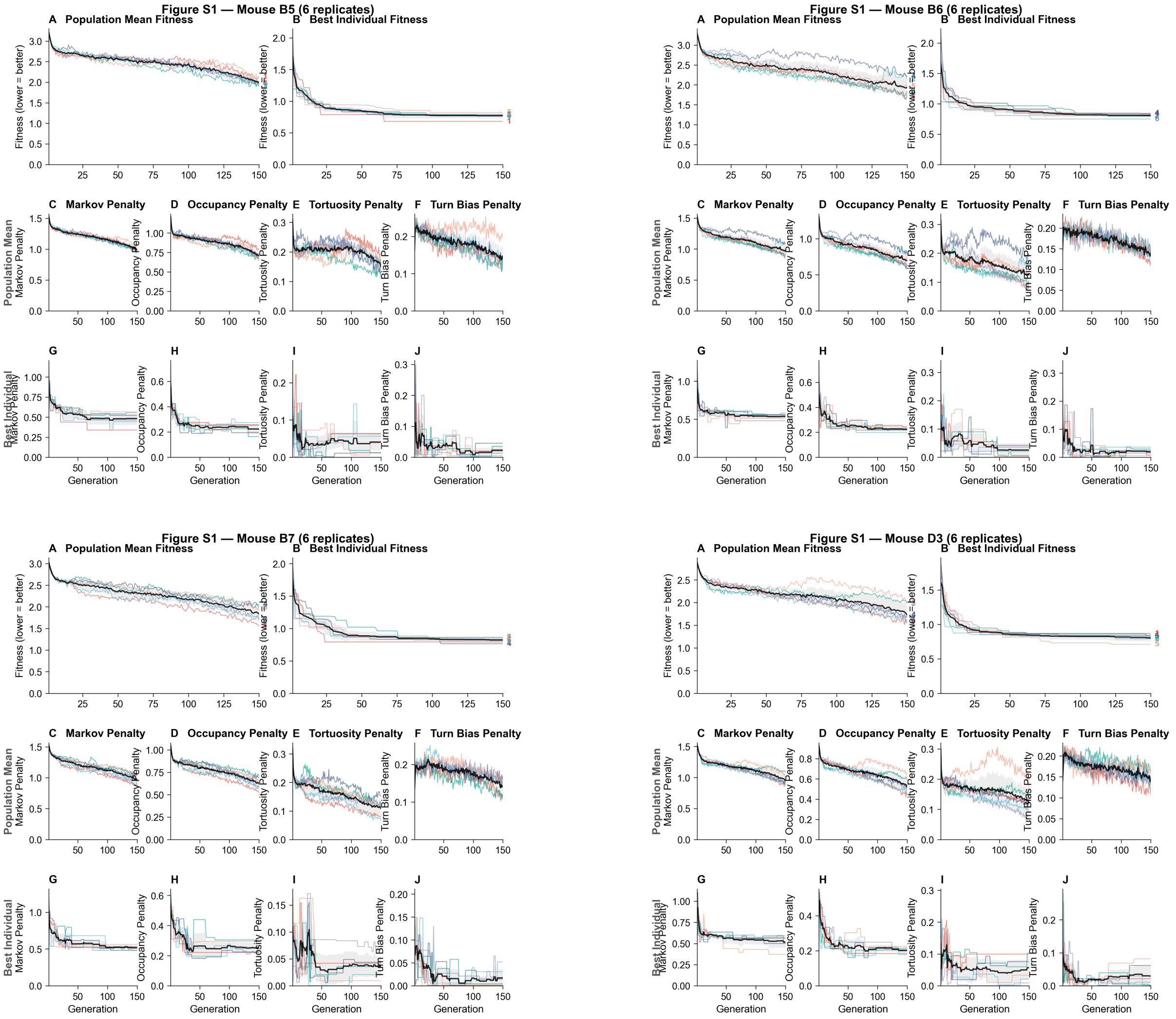


**Figure S1. Per-mouse individual replicate trajectories (mice B5–D3).** Population mean fitness (solid lines) and best-of-generation fitness (dashed lines) across 150 generations. Lower fitness = better behavioral match to the target mouse. Continued in the next part for the remaining five mice.

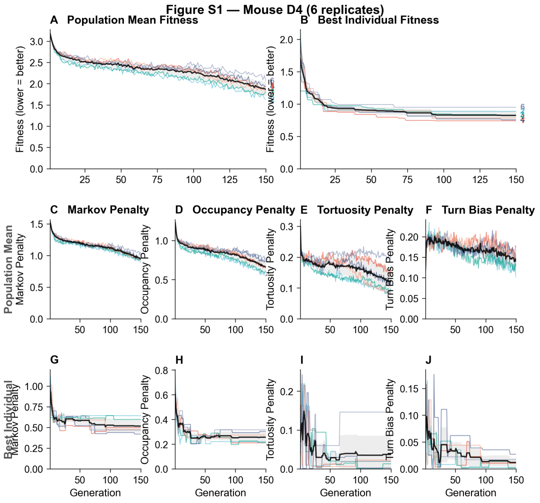


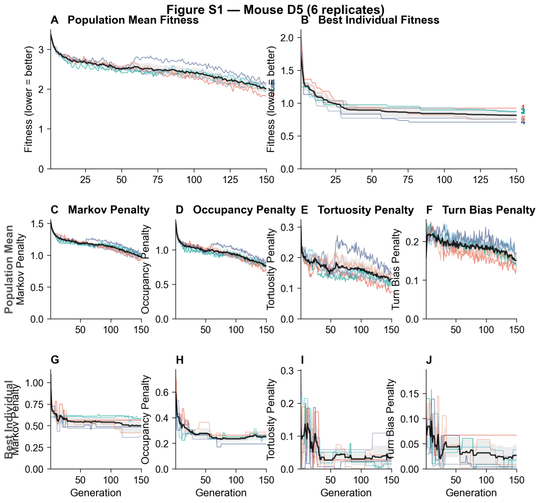


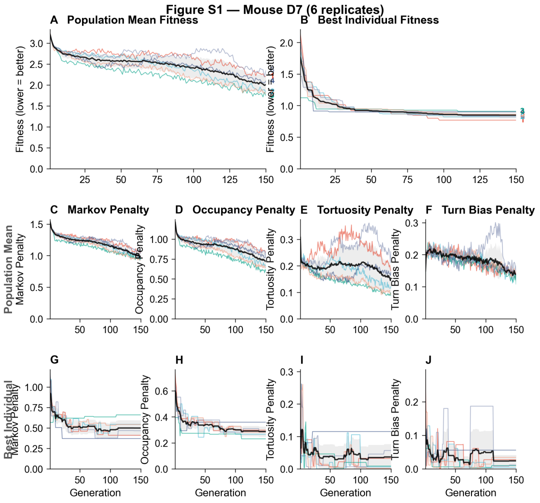


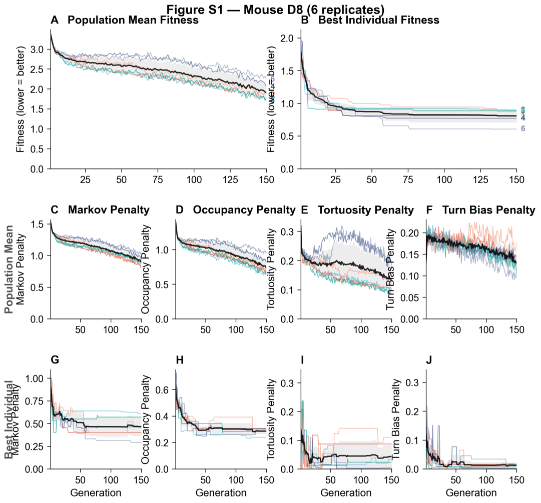


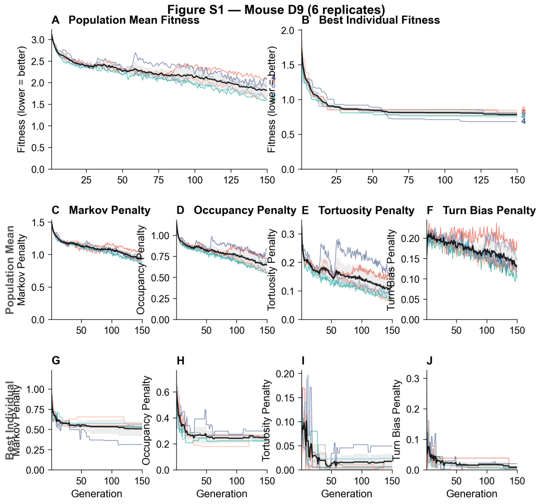


**Figure S1. Per-mouse individual replicate trajectories (mice D4–D9, continued).**

#### S2. Evolved network diagrams per mouse

Representative best-evolved network diagram for each of the 9 mice. Excitatory connections are shown in red, inhibitory in blue; connection width encodes weight magnitude. Despite identical aggregate structural statistics, the specific who-connects-to-whom topology varies across mice and replicates.

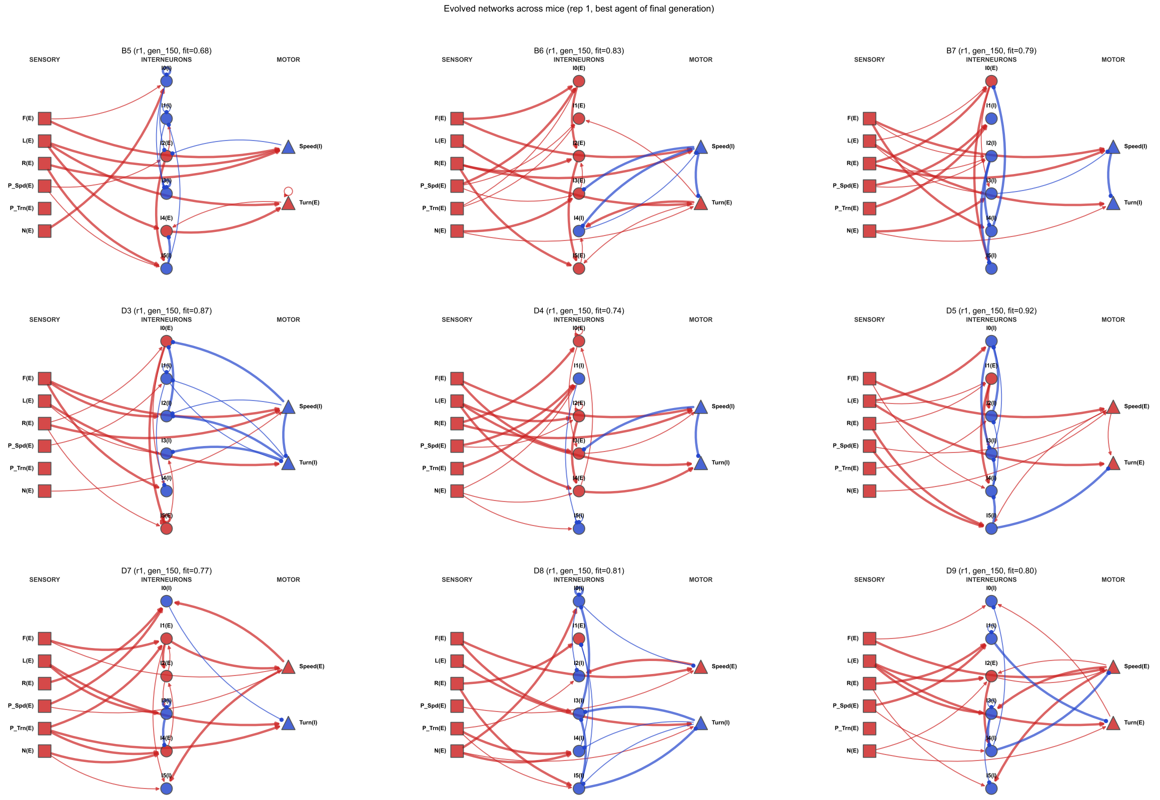


**Figure S2. Evolved network diagrams for all 9 mice.** Best-evolved agent per mouse at generation 150. Node types: squares = sensory, circles = interneuron, triangles = motor. Red = excitatory, blue = inhibitory; line width ∝ weight magnitude.

#### S3. Per-metric convergence curves

Figure S3 shows the evolutionary convergence broken down by the four fitness components, with population mean and best-of-generation curves per mouse. KDE insets (panels C–F) show the distribution of generation-150 fitness values across replicates for each mouse. The Markov and occupancy components (weighted 2×) account for most early-generation improvement.

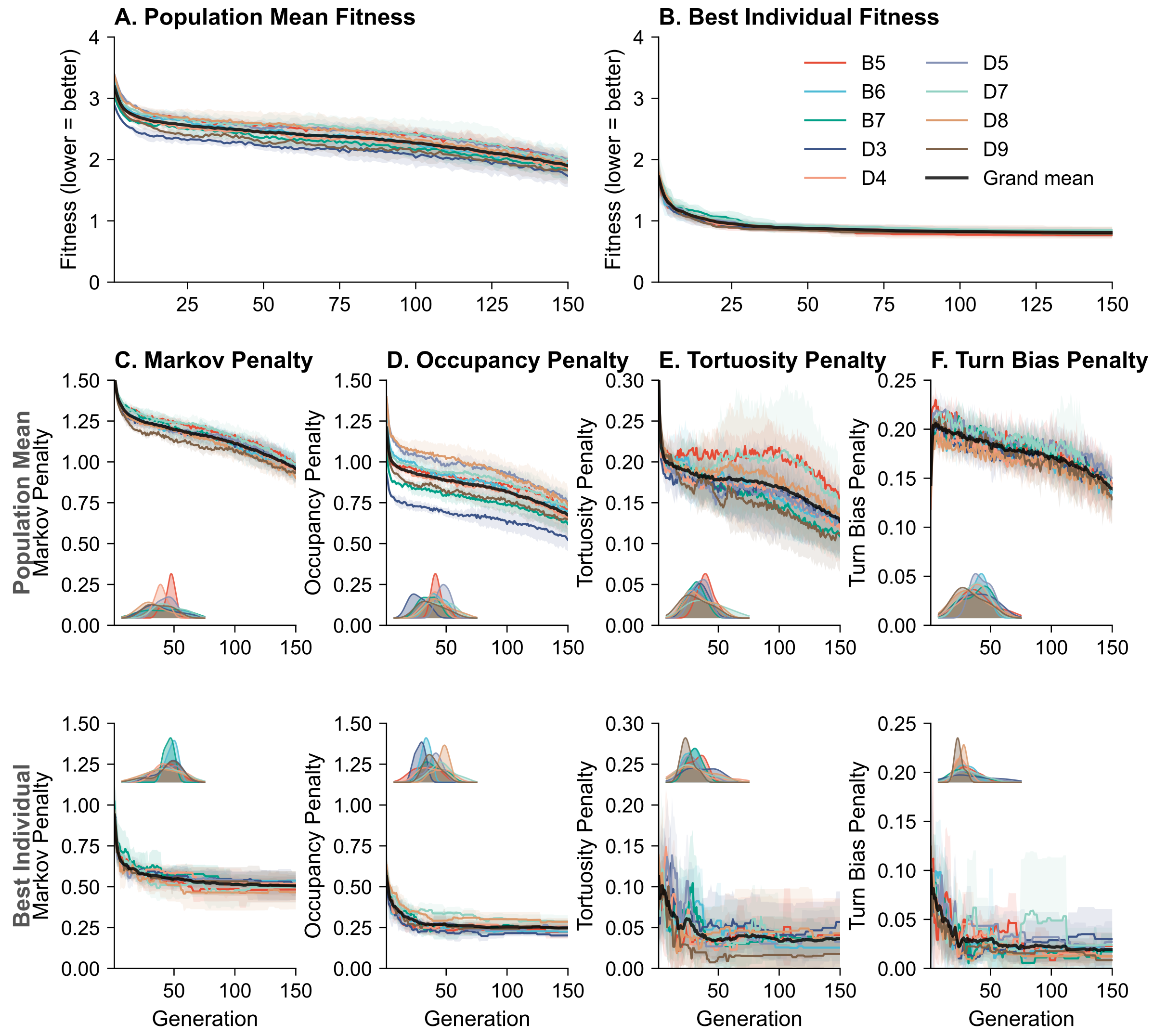


**Figure S3. Per-metric evolutionary convergence across 9 mice.** (A) Population mean total fitness (all mice, 6 replicates each). (B) Best individual total fitness. (C–F) Population mean fitness (top sub-row) and best-of-generation fitness (bottom sub-row) for each of the four behavioral components: Markov (C), Occupancy (D), Tortuosity (E), Turn Bias (F). KDE insets show the generation-150 fitness distribution per mouse. Lower = better match.

#### S3b. Evolved vs random circuit features (numerical summary)

Table S1 lists the seven key structural features from Figure 1G with exact means, Cohen's $d$, and $p$-values for reference. The forest plot (Figure 1G) shows all 18 features with 95% bootstrap CIs.

**Table S1. Evolved vs random circuit features (selected).** One-sample $t$-test (evolved values vs random mean; $n_{\text{evolved}} = 54$, $n_{\text{random}} = 200$). Cohen's $d$ measures effect size (positive = evolved > random). Full 18-feature comparison with bootstrap CIs is shown in Figure 1G.

| Feature | Evolved | Random | Cohen's $d$ | $p$ |
|---|---|---|---|---|
| Total connections | 23.426 | 24.445 | −1.10 | $9.3 \times 10^{-16}$ |
| Sensory-to-motor shortcuts | 3.537 | 2.470 | +1.11 | $9.2 \times 10^{-16}$ |
| Inter-to-motor connections | 1.759 | 2.685 | −0.93 | $2.0 \times 10^{-10}$ |
| Motor-to-motor connections | 0.667 | 0.975 | −0.41 | 0.0009 |
| Mean weight magnitude | 0.670 | 0.621 | +0.74 | $4.1 \times 10^{-7}$ |
| Fraction at max weight | 56% | 49% | +0.74 | $4.1 \times 10^{-7}$ |
| Interneuron mean fan-in | 2.910 | 3.053 | −0.95 | $6.6 \times 10^{-13}$ |

#### S4. Emergent behavioral properties: comparison to real mice

Thigmotaxis (wall contact fraction), median speed, and mean absolute turn rate were computed for all 54 best-evolved agents and compared to their target mouse. Each property was reproduced without explicit selection: evolved agents were not optimised for wall proximity, speed magnitude, or turn rate, yet all three statistics fall within the range of real mouse values across all 9 mice (Figure S4). Speed and turn rate distribution panels appear in Figure 1F–G in the main text.

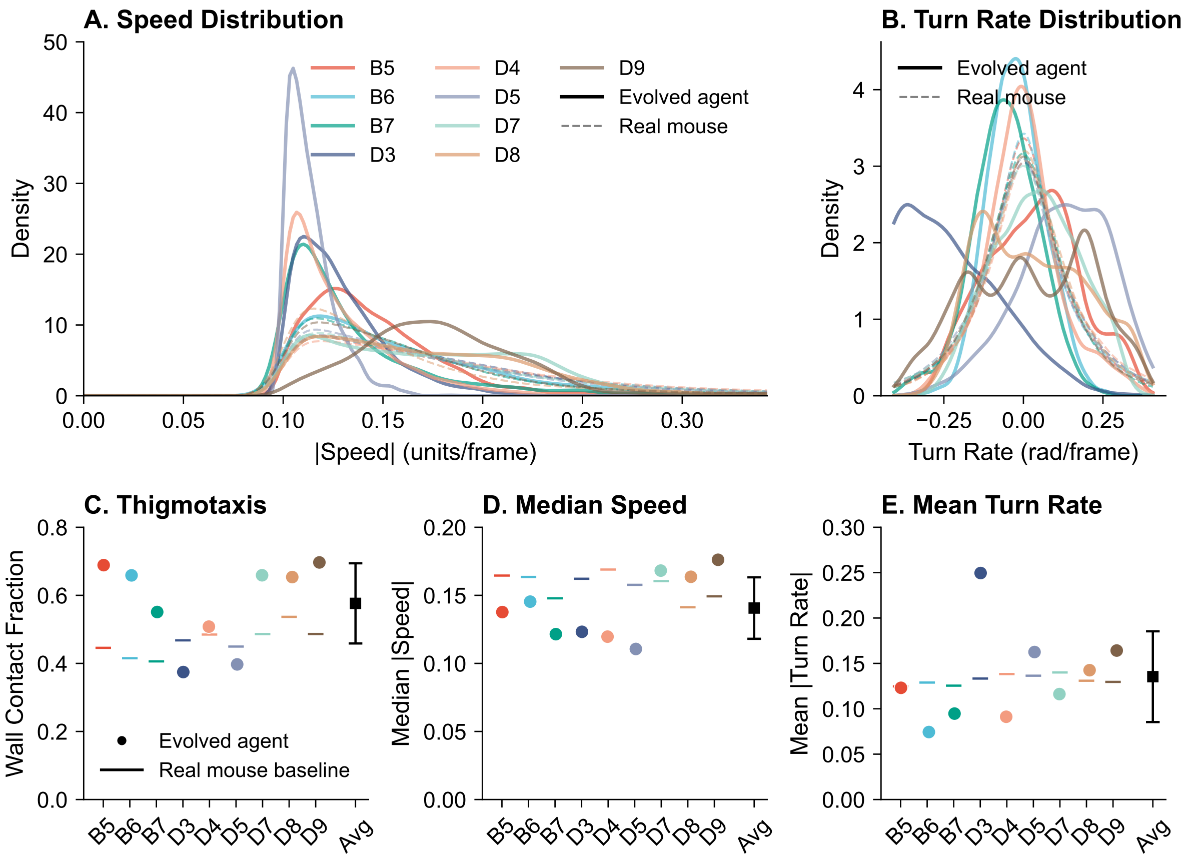


**Figure S4. Emergent behavioral properties per mouse (panels C–E).** Panels A–B (speed and turn rate distributions) appear in Figure 1F–G. (C) Wall contact fraction (thigmotaxis): strip plots for evolved replicates (dots) overlaid on real mouse baseline (horizontal line). (D) Median speed. (E) Mean absolute turn rate. In all panels, evolved agent values bracket the real mouse value, confirming that these emergent properties arise from the general behavioral policy.

#### S5. Excitatory-inhibitory balance dynamics

Figure S5 shows the temporal evolution of E/I balance across generations for interneurons and motor neurons. Interneurons show no consistent convergence pattern (panel A). The speed motor neuron converges uniformly to a strongly excitatory-dominated regime (mean $\approx$ 0.85) across all 9 mice (panel B). The turn motor neuron converges to a more balanced regime (mean $\approx$ 0.64) with greater inter-mouse variability (panel C). The per-neuron E/I heatmap at generation 150 appears in Figure 1H.

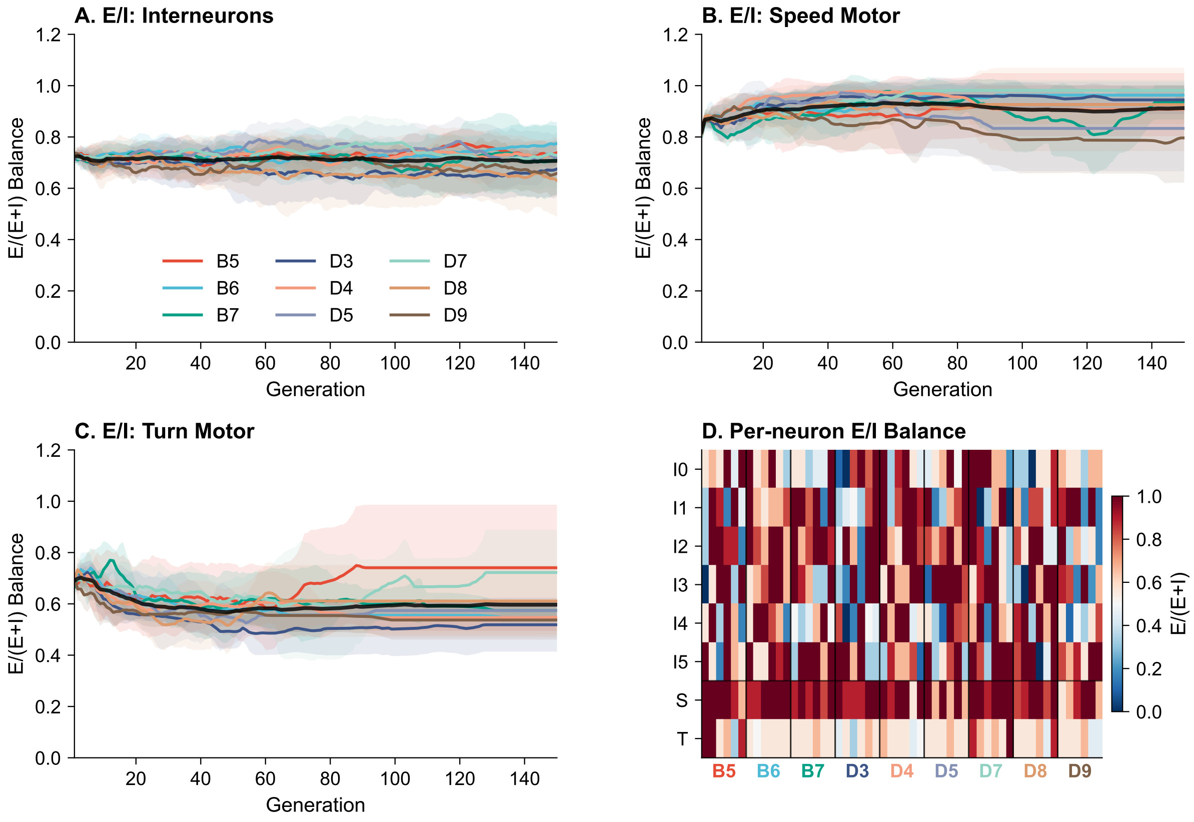


**Figure S5. Excitatory-inhibitory balance dynamics (panels A–C).** Panel D (per-neuron E/I heatmap at generation 150) appears in Figure 1H. (A) E/I balance for all six interneurons over 150 generations (coloured lines = per-mouse mean ±SD; black = grand mean). No consistent convergence pattern emerges. (B) E/I balance for the speed motor neuron, converging uniformly to mean ≈ 0.85 excitatory across all 9 mice. (C) E/I balance for the turn motor neuron, converging to mean ≈ 0.64 with greater variability, consistent with closed-loop inhibitory modulation of turning direction.

#### S6. Evolved vs random weight similarity distributions

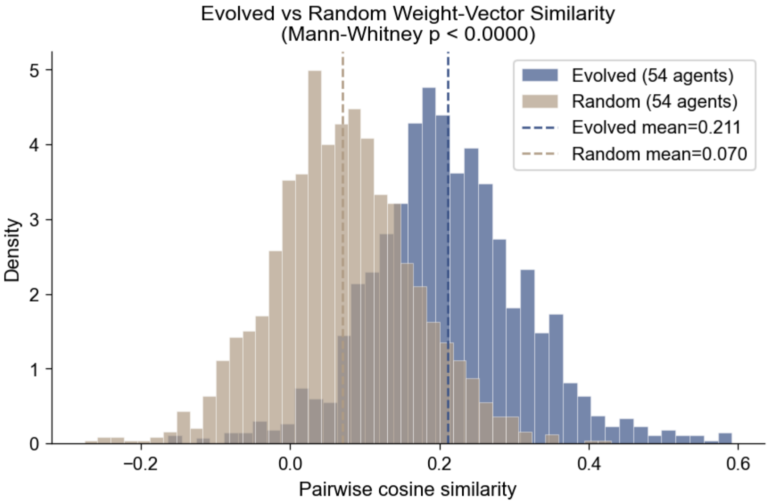


**Figure S6. Evolved circuits occupy a significantly narrower region of weight space than random agents.** Pairwise cosine similarity distributions for evolved (blue; $n = 54$) and randomly initialised constrained agents (tan; $n = 200$). Evolved mean $= 0.211$; random mean $= 0.070$; Mann-Whitney $p = 1.2 \times 10^{-229}$. Evolution compresses all circuits into a shared structural region, though the $54 \times 54$ similarity matrix (Figure 2A) shows no mouse-level block structure within that region.

#### S7. Power analysis

Post-hoc power analysis for the 18-feature ANOVA is shown in Figure S7. At $N = 54$ and $k = 9$ groups, the minimum detectable $\eta^2$ at 80% power is 0.247. The three largest-effect features each exceed this threshold (82–88% power), confirming the null result is not due to insufficient power for large effects. Medium-effect features ($\eta^2 \approx 0.06$–$0.14$) would require 12–28 replicates per mouse.

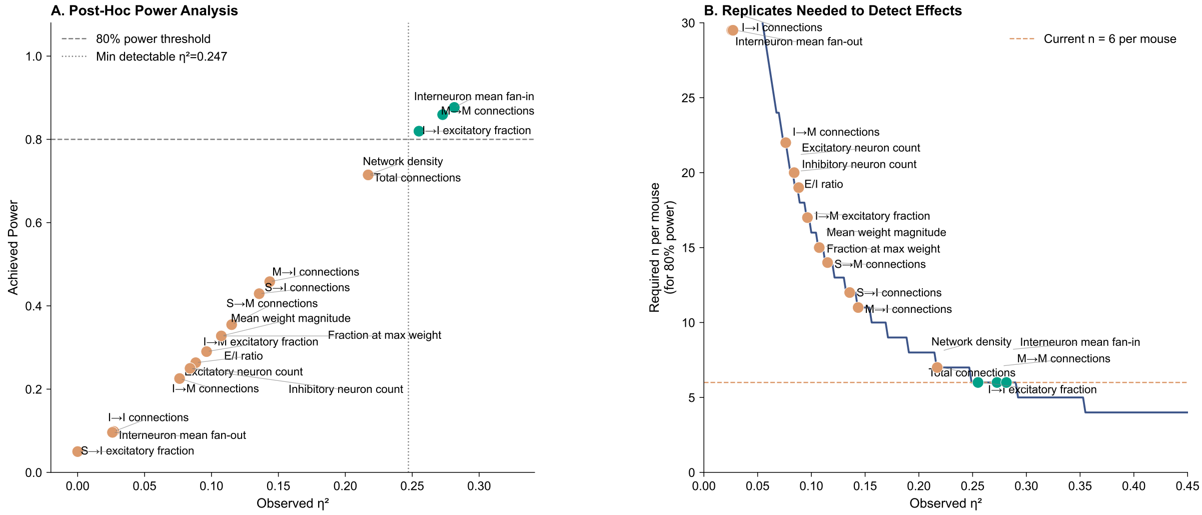


**Figure S7. Post-hoc power analysis.** (A) Observed $\eta^2$ vs achieved power. Teal points exceed 80% power threshold; orange points fall below it. (B) Required replicates per mouse for 80% power at each observed effect size. Current $n = 6$ per mouse (dashed orange line) is sufficient for large effects.

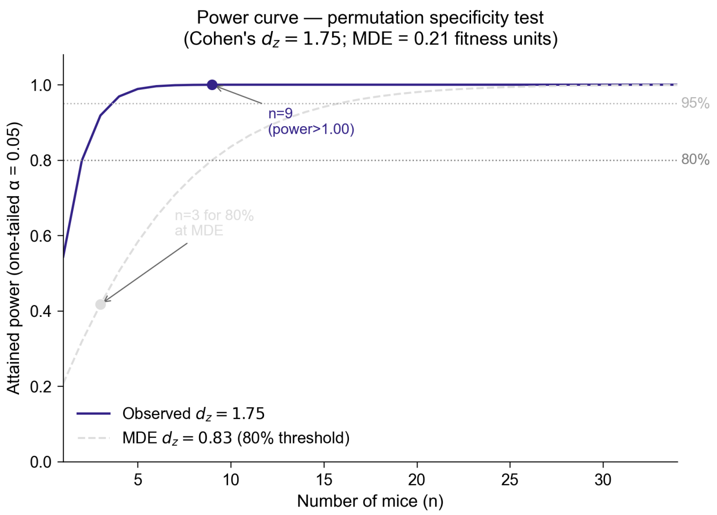


**Figure S8. Power curve.** Required replicates per mouse for 80% power across a range of effect sizes ($\eta^2$).

#### S8. Per-metric specialisation: cross-evaluation heatmaps

The cross-evaluation structure visible in the total fitness matrix (Figure 2D) replicates independently within each fitness component. Each of the four per-metric $9 \times 9$ matrices shows the same diagonal pattern: agents consistently achieve lower error on their own target mouse than on other mice (Figure S9).

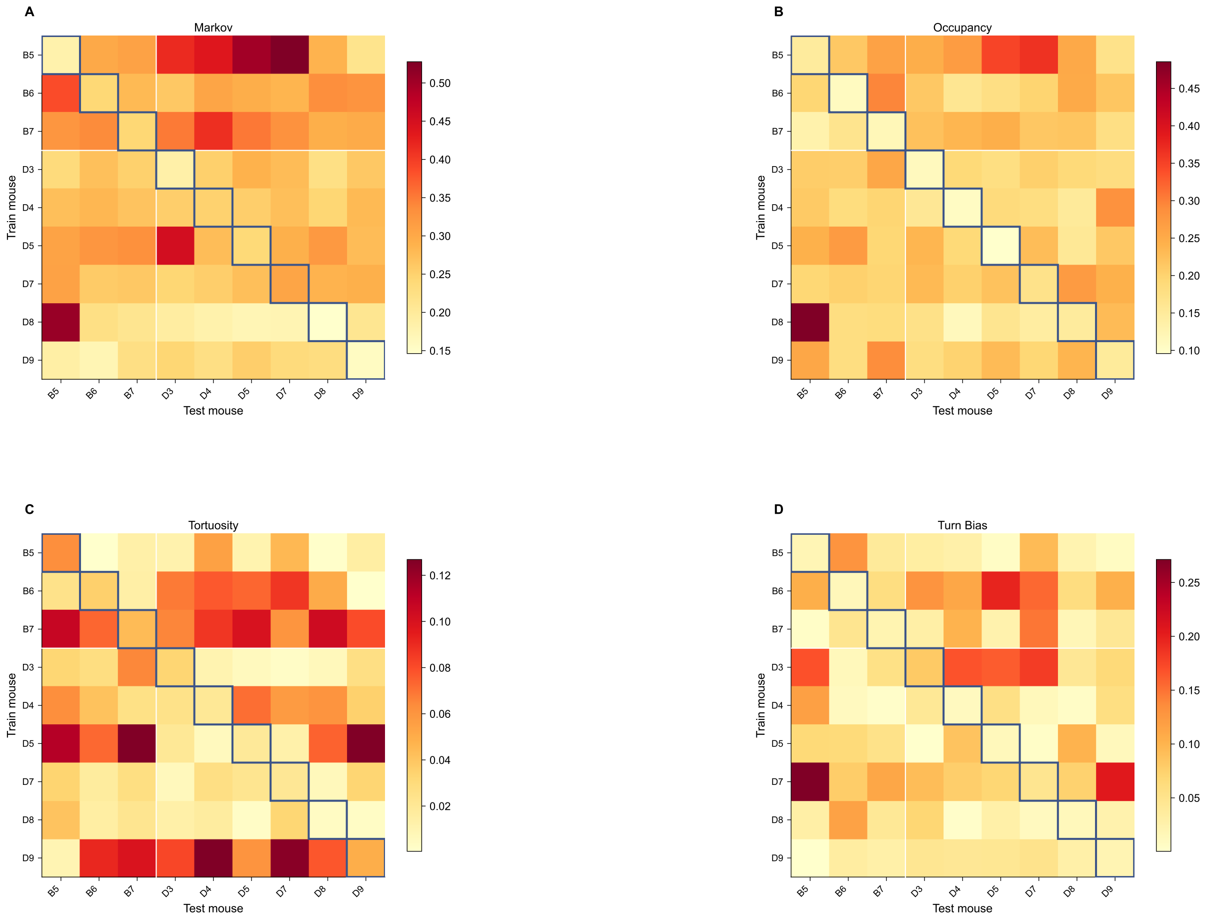


**Figure S9. Per-metric** $9 \times 9$ **cross-evaluation matrices.** Each panel shows the cross-evaluation fitness matrix for one of the four behavioural components (Markov, Occupancy, Tortuosity, Turn Bias). Colour encodes component fitness error (lower = better match). Blue boxes mark diagonal (own-mouse) entries. The diagonal specialisation structure replicates independently in every component.

#### S9. Per-metric specialisation: own vs cross-mouse bars

Cross-mouse error exceeds own-mouse error in every fitness component; the own/cross ratio ranges from 0.400 (turn bias, most individual-specific) to 0.749 (Markov transitions; Figure S10).

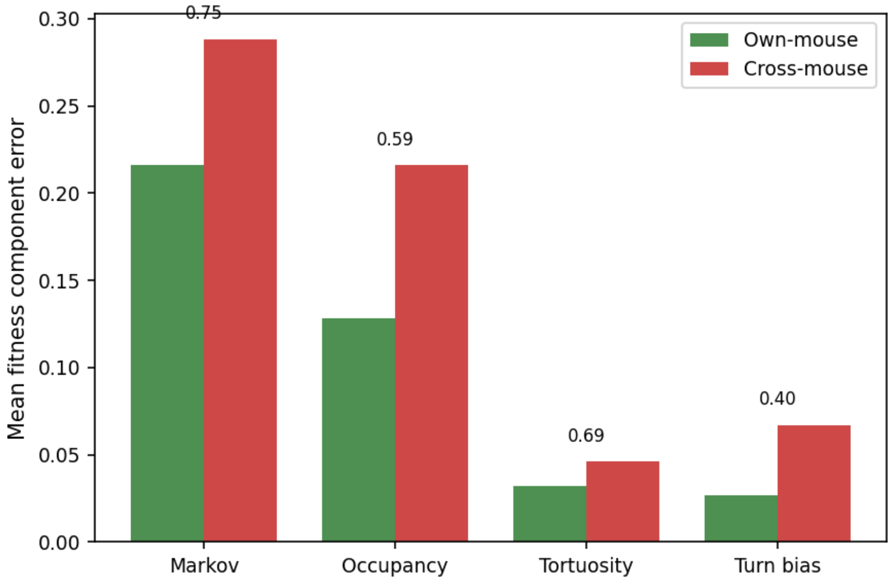


**Figure S10. Own-mouse vs cross-mouse error per fitness component.** Mean fitness component error for own-mouse (green) and cross-mouse (red) evaluations, averaged across all 9 mice. Ratio annotated above each cross-mouse bar (own/cross; lower = more specialised). Turn bias shows the strongest individual-specificity (ratio = 0.400), followed by occupancy (0.594), tortuosity (0.693), and Markov transitions (0.749).

#### S10. Strain control

Specialisation indices computed within each genetic strain independently (B-strain: 0.648; D-strain: 0.734) remain close to the overall index (0.334), confirming that individual-level behavioral divergence is present within each strain and is not an artefact of between-strain differences (Figure S11).

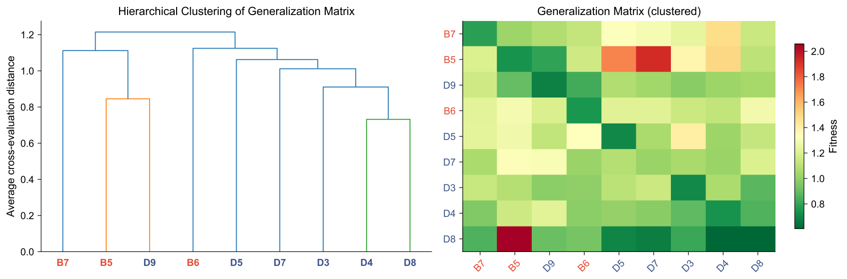


**Figure S11. Strain control.** Specialisation indices within B-strain and D-strain mice remain positive and close to the overall index.

#### S11. Fitness weighting scheme robustness

The specialisation index was recomputed under five alternative fitness weighting schemes for the four behavioral components (Markov, Occupancy, Tortuosity, Turn Bias). All five schemes produce positive mean indices (Table S2; Figure S12), confirming the finding is not sensitive to the choice of component weights.

**Table S2. Specialisation index under five fitness weighting schemes.** The primary analysis uses the Published 2:2:1:1 scheme. Per-mouse breakdown is shown in Figure S12.

| Weighting scheme | Mean specialisation index | Mice > 0 |
|---|---|---|
| Published (2:2:1:1) | 0.334 | 9/9 |
| Equal (1:1:1:1) | 0.347 | 9/9 |
| Turn-dominant (1:1:1:3) | 0.392 | 9/9 |
| Markov-only (1:0:0:0) | 0.251 | 8/9 |
| Turn+Occupancy (0:2:0:1) | 0.432 | 9/9 |

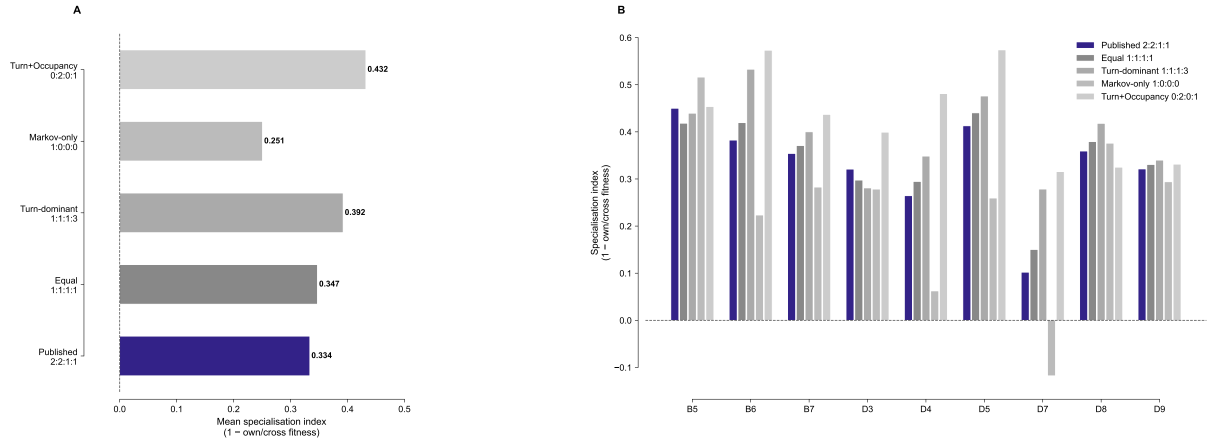


**Figure S12. Specialisation index is robust to fitness weighting choice.** (A) Mean specialisation index (1 − own/cross fitness) across all 9 mice for each of five weighting schemes. Published 2:2:1:1 (highlighted) is the primary analysis; dashed line at zero indicates no specialisation. (B) Per-mouse specialisation index for all 9 mice across the five schemes. All schemes produce positive mean indices.

#### S12. Specialisation persists on held-out behavioral data

Behavioral baseline and fitness evaluation used the same trajectory bouts from each mouse's recording. To test whether specialisation reflects genuine individual-level encoding rather than bout-specific fitting, we performed cross-bout evaluation using the second half of each mouse's trajectory as a held-out test set.

The held-out specialisation index was 0.096 across all 9 mice (all held-out indices $> 0$; Figure S13), compared to 0.334 on the training set. The attenuation reflects reduced statistical precision of the held-out behavioral baseline rather than overfitting: each mouse's held-out target is estimated from ≈99 trajectory bouts rather than ≈197, reducing the resolution of the behavioral fingerprint. No agent lost specialisation entirely; the unanimous persistence of positive indices is inconsistent with genuine overfitting.

We confirmed directly, from the real mouse data alone, that the individual behavioral targets are stably distinguishable beyond within-mouse bout-to-bout noise (this is the foundation on which the agent-level specialisation rests). We partitioned each mouse's bouts into 6 folds, computed the four fitness metrics per fold with the same machinery used for fitness, and compared within- vs between-mouse fold distances. Within-mouse fold distances were smaller than between-mouse distances in the combined fitness metric (within = 0.448, between

$= 0.629$; Mann-Whitney $p < 0.001$, rank-biserial $= 0.694$), and a leave-one-fold-out nearest-mouse classifier reached 87% accuracy against a 11% chance level (permutation $p < 0.001$). The mice are therefore genuinely individuated in the behavioral metrics, so the held-out attenuation is target-estimation noise, not a circularity of the fitness definition.

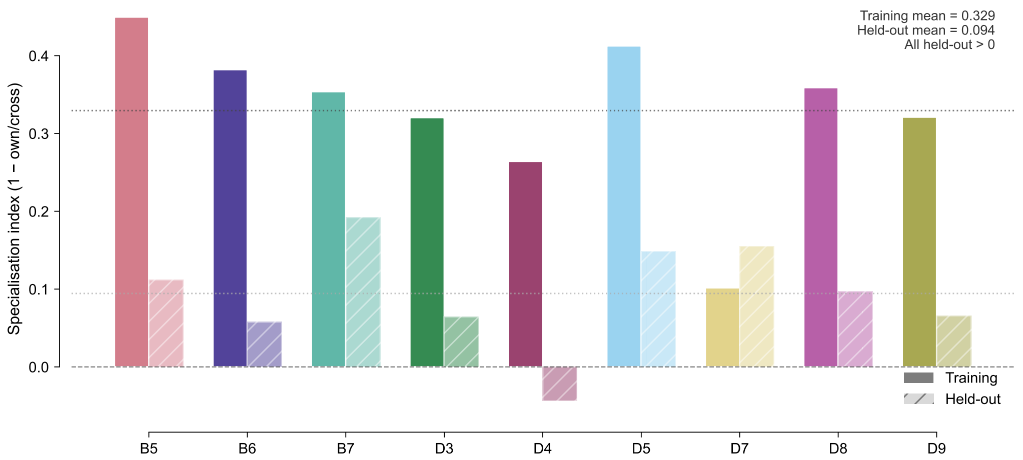


**Figure S13. Holdout specialisation indices per mouse.** Paired bars showing training-set (solid) and held-out (hatched) specialisation indices for each of the 9 mice, coloured by mouse identity. All held-out indices are positive (population mean $= 0.096$), confirming that specialisation generalises across trajectory bouts. The attenuation relative to training (0.334) reflects reduced bout count in the held-out half, not overfitting.

#### S12b. Per-source ablation sensitivity (null result)

Permuting each source neuron's outgoing connections individually, rather than the full topology at once, yields a $54 \times 14$ sensitivity matrix (Figure S14). Agents are ordered by mouse (six replicates per mouse, separated by white lines). No mouse-level block structure is visible: 0 of 14 source neurons reach significance after Bonferroni correction, and within-mouse sensitivity profiles are no more correlated than between-mouse profiles ($r_{\text{within}} = 0.150$, $r_{\text{between}} = 0.167$, Mann-Whitney $p = 0.6919$). Individual source neurons do not fingerprint mouse identity, even though the full topology is causally necessary.

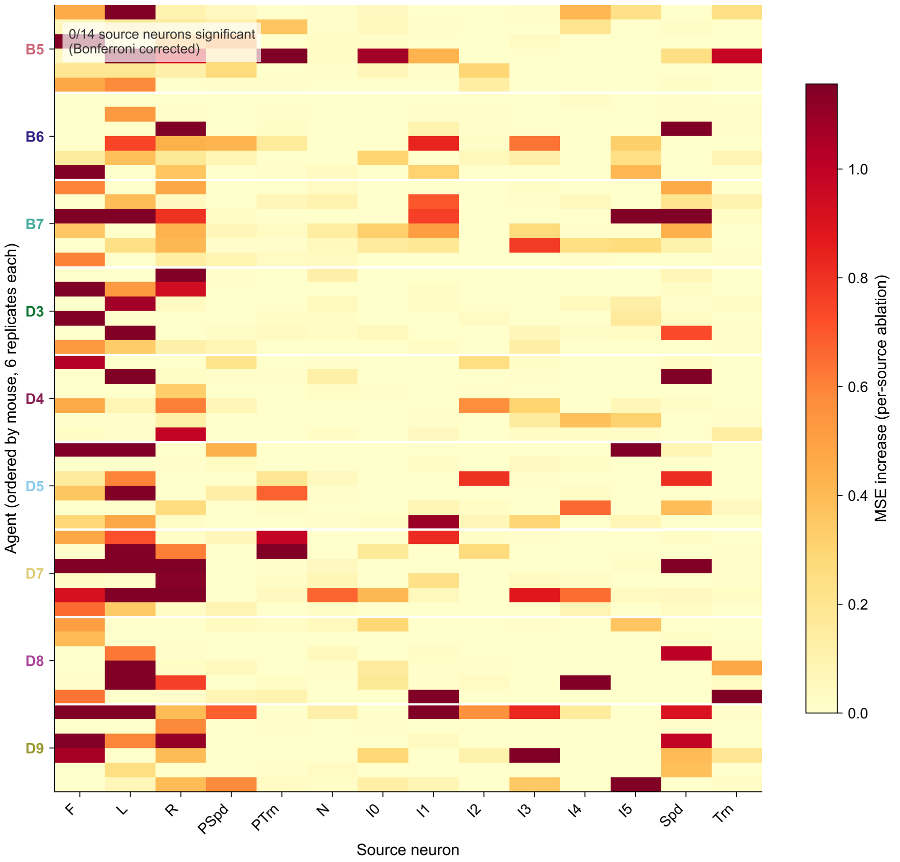


**Figure S14. Per-source ablation sensitivity matrix (54 agents $\times$ 14 source neurons).** Each cell shows the increase in motor output MSE when that source neuron's outgoing connections are permuted individually. Agents are ordered by mouse (white separator lines every 6 rows). No mouse-level block structure is visible; 0/14 source neurons reach significance after Bonferroni correction.

#### S13. Structural clustering visualization: k-means groups do not correspond to mouse identity or performance

The mutual information analysis of Section 2.4 groups each of the 54 agents into one of $k = 5$ structural clusters via k-means, separately for each structural axis, and measures how well these cluster labels predict mouse identity. Figure S15 shows two complementary views of why these structural clusters carry no behavioral information.

Panels A and B show the same PCA embedding of agents in topology feature space with different color encodings. Panel A colors by k-means cluster: five structural groups are present in the data. Panel B colors the same points by mouse identity: agents from the same mouse are scattered across all five clusters, and each cluster contains agents from multiple mice. Panel C replicates the null on the magnitude axis: mouse identity is equally unrecoverable from structural similarity in magnitude space. Panel D provides the fitness-quintile companion analysis: agents ranked by own-mouse fitness (best on left), colored by topology cluster. If structurally similar agents performed similarly, colors would grade smoothly across the rank axis; they do not.

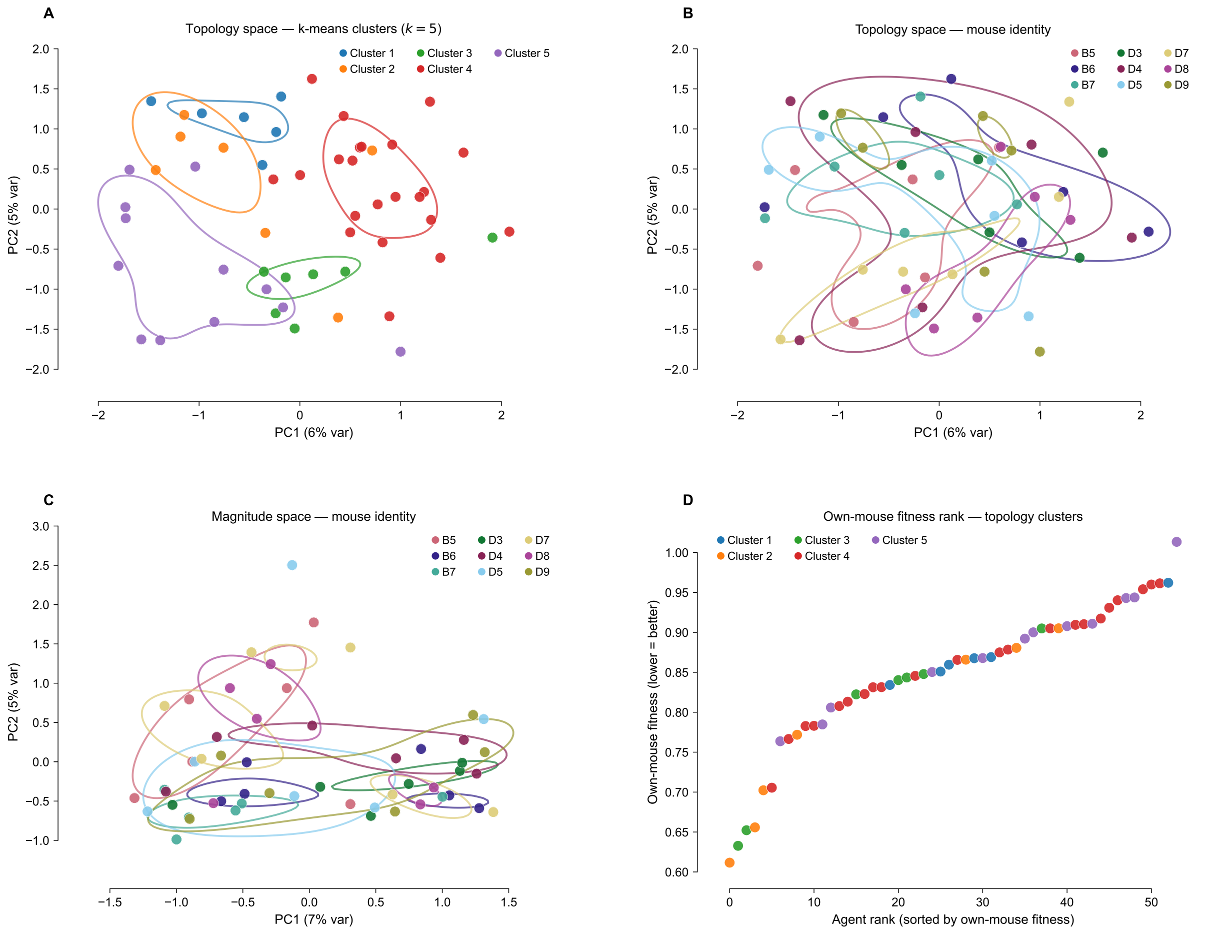


**Figure S15. Structural k-means clusters do not correspond to mouse identity or behavioral performance.** (A) PCA of all 54 agents in topology feature space ($14 \times 14$ binary adjacency, 196-dimensional), colored by k-means cluster assignment ($k = 5$, blue palette). Five structural groups are visible. (B) Same topology PCA, colored by mouse identity (Tol palette, 9 mice). Agents from the same mouse are distributed across all structural clusters: topology is degenerate with respect to mouse identity. (C) PCA of magnitude feature space, colored by mouse identity. The null replicates on a different structural axis. (D) All 54 agents ranked by own-mouse fitness (lower = better; best on left), colored by topology cluster. Mixed colors across the performance rank confirm that structurally similar agents do not perform similarly: the fitness-quintile companion to the mouse-identity MI analysis in the main text. Percentage variance explained by PC1 and PC2 shown on axes A–C.

#### S14. Sensitivity RSA: the geometry of sensitivity profiles does not fingerprint mouse identity

To test whether the full 14-dimensional sensitivity profile holistically fingerprints mouse identity, we performed a representational similarity analysis (RSA). We computed pairwise Euclidean distances between agents' 14-dimensional ablation sensitivity vectors, yielding a $54 \times 54$ sensitivity RSM. We then tested whether within-mouse pairs have lower sensitivity distance than between-mouse pairs (Mann-Whitney $U$, $p = 0.529$) and whether the sensitivity RSM correlates with a mouse-identity RSM (Mantel test: $\rho = -0.044$, permutation $p = 0.357$; 10,000 permutations).

Both tests are null. The *geometry* of the sensitivity profile does not fingerprint mouse identity: agents trained on the same mouse are no more similar in their sensitivity profiles than agents trained on different mice. This is consistent with the per-source ablation null (0/14 neurons significant; reported in Section 2.5), and together they rule out any structural decomposition of the sensitivity profile as a mouse-identity fingerprint. The result that individuates specialists from generalists is not the *pattern* of sensitivity but its *variance* (Section 2.6).

Note: this analysis is likely underpowered at $n = 6$ replicates per mouse. Additional replicates are planned (see Section 3.4).

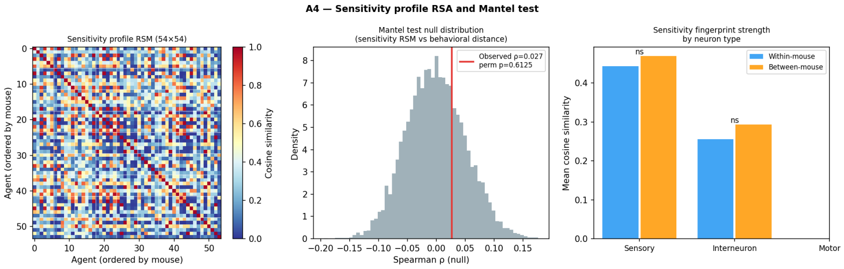


**Figure S16. Sensitivity RSA: no mouse-level structure.** Left: $54 \times 54$ sensitivity RSM (pairwise Euclidean distance between ablation sensitivity vectors), agents ordered by mouse. No block-diagonal structure is visible. Right: within-mouse vs between-mouse sensitivity distance distributions (Mann-Whitney $p = 0.529$).

#### S15. Representational geometry: positive control and robustness

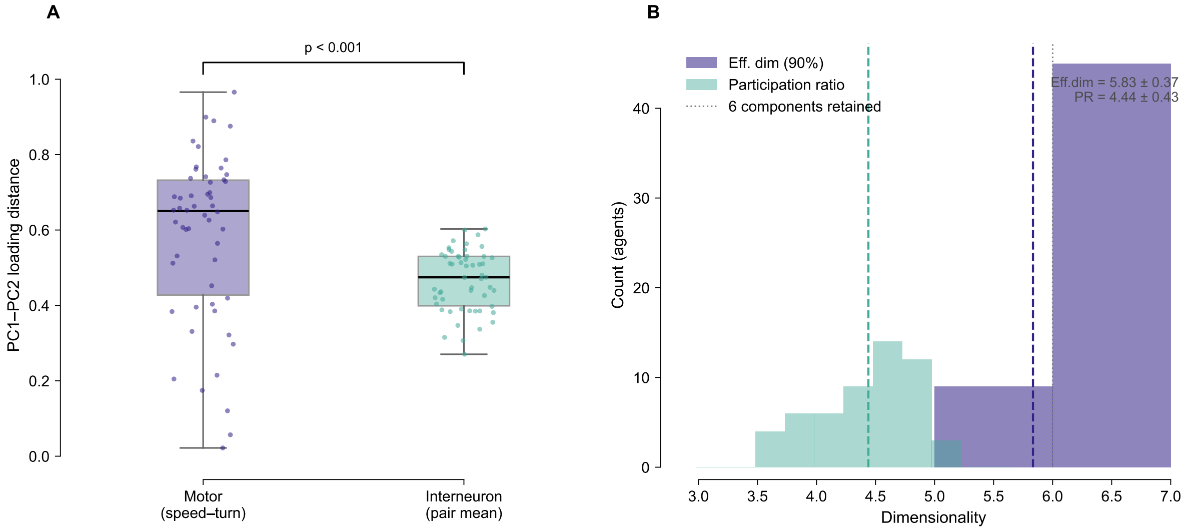


**Figure S17. Activity embedding positive control.** Motor neuron separation in PC1–PC2 loading space as a sanity check for embedding informativeness at the small scale of these networks ($N = 14$). Speed (neuron 12) and Turn (neuron 13) motor neurons are consistently more separated from each other than the typical interneuron pair ($0.583 \pm 0.220$ vs $0.466 \pm 0.080$; Mann-Whitney $p < 0.001$), confirming that neuronal embeddings capture functional circuit structure despite the small network size.

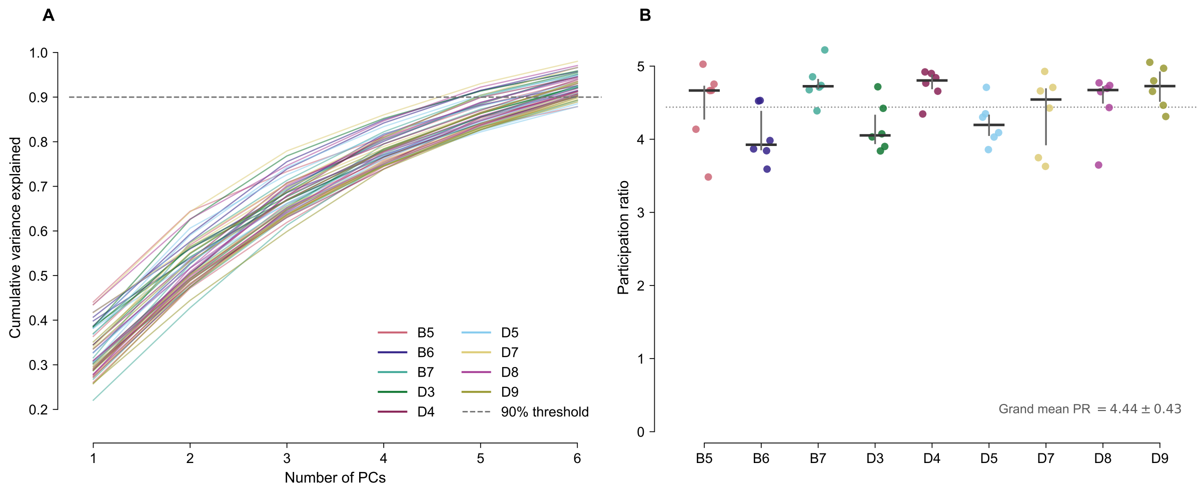


**Figure S18. Manifold dimensionality is near-maximal and uniform across mice.** *Left:* Cumulative variance explained by the top 6 PCs for each of the 54 best-evolved agents (lines coloured by mouse). The 90% threshold (dashed grey) is reached by PC 5–6 in all agents, justifying retention of 6 components. Curves are highly consistent across mice, with no mouse-specific variance structure. *Right:* Participation ratio (PR) per mouse (strip plot with median and IQR; one dot per agent). Grand mean PR $= 4.44 \pm 0.43$ out of a maximum of 6, corresponding to effective dimensionality $5.83 \pm 0.37$. No mouse shows a distinctively lower or higher dimensionality, confirming that individual-target selection does not compress or expand the dimensionality of representational space.

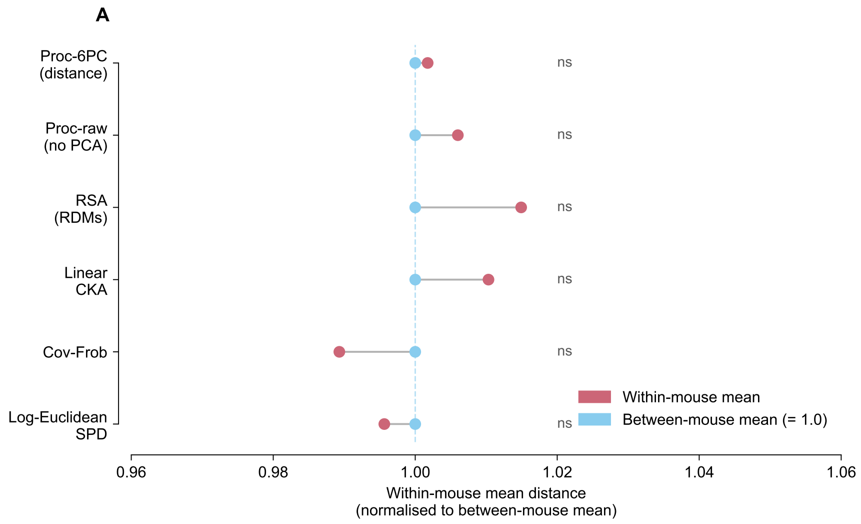


**Figure S19. Representational degeneracy is robust across six independent similarity metrics.** Within-mouse mean distance (red) versus between-mouse mean distance (blue, set to 1.0 for each method) for six metrics applied to the same 54 agent activity records. Proc-6PC: Procrustes distance on 6-PC loading matrices (main analysis). Proc-raw: Procrustes on raw per-bout means without PCA. RSA: representational dissimilarity analysis on per-bout response RDMs. CKA: centred kernel alignment. Cov-Frob: Frobenius distance between neuron-neuron covariance matrices. Log-Euclidean SPD: affine-invariant distance on the SPD manifold. All pairwise comparisons are indistinguishable (all MW $p > 0.25$, ns), ruling out PCA truncation or a particular distance convention as an explanation for the null result.

#### S16. Dynamical null results

Trajectory RSM and Lyapunov analyses used a standardised synthetic sensory input sequence (Section 3.13). Activation trajectory cosine similarities and Lyapunov exponents ($\lambda_1$) showed no between-mouse differences (trajectory RSM: $p = 0.5085$; Lyapunov: $p = 0.943$; fixed-point eigenvalue: $p = 0.460$). Grand mean $\lambda_1 = -0.050 \pm 0.024$, confirming a uniform slightly stable dynamical regime across all mice.

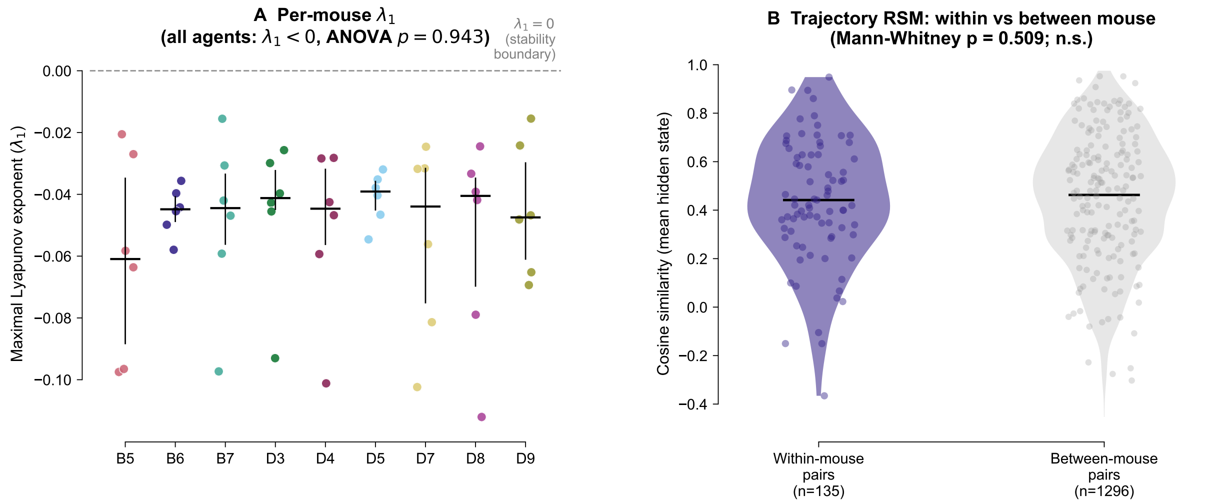


**Figure S20. Dynamical null results.** (A) Per-mouse maximal Lyapunov exponent $\lambda_1$ (strip plot; median and IQR shown). All 54 agents are slightly stable ($\lambda_1 < 0$); one-way ANOVA returns $F = 0.345$, $p = 0.943$ — no mouse-specific stability signature. (B) Within-mouse (purple) versus between-mouse (grey) activation trajectory cosine similarity (violin + strip). Distributions are indistinguishable (Mann-Whitney $p = 0.5085$; n.s.).

#### S17. Attractor landscape analysis

Each agent was run from 200 random initial states with zero sensory input for 500 steps to map its autonomous attractor landscape. Final interneuron and motor states (8-dimensional) define an attractor fingerprint per agent. Pairwise sliced Wasserstein distances between agent attractor distributions showed no within-mouse clustering (within $0.081 \pm 0.125$ versus between $0.079 \pm 0.124$; Mann-Whitney $p = 0.544$).

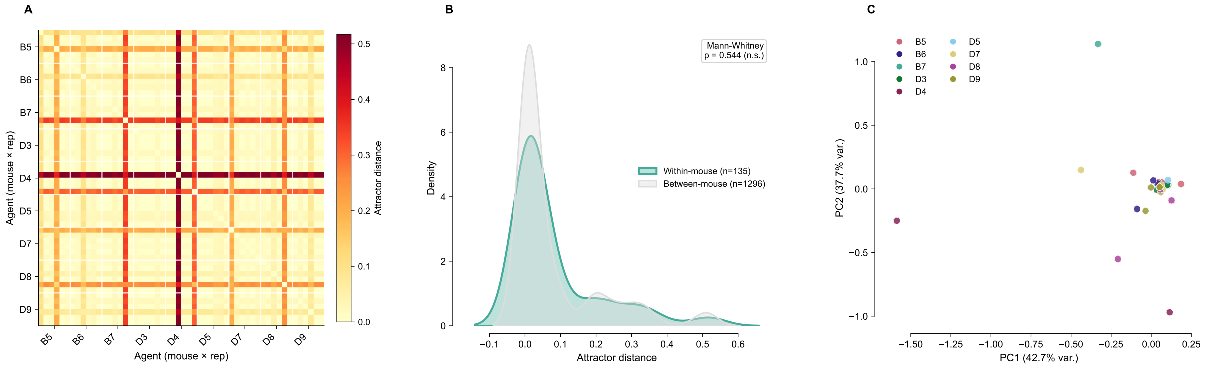


**Figure S21. Attractor landscape is degenerate across mice.** *Left:* $54 \times 54$ sliced Wasserstein distance matrix between agent attractor distributions (plasma scale). No block structure is visible. *Centre:* KDE of within-mouse (red) versus between-mouse (blue) attractor distances (Mann-Whitney $p = 0.544$; n.s.). *Right:* Attractor fingerprints in 2D (PCA of zero-input final states; one representative agent per mouse, 200 initial conditions each). Points coloured by mouse. No mouse-specific clustering is apparent.

Per-mouse specialisation trajectories. Figure S22 shows the specialisation index (mean ±

SD across 6 replicates) for each of the 9 mice independently over 150 generations. All mice develop positive specialisation indices; the rate of emergence and final magnitude vary across mice, consistent with the between-mouse variance visible in Figure 2E.

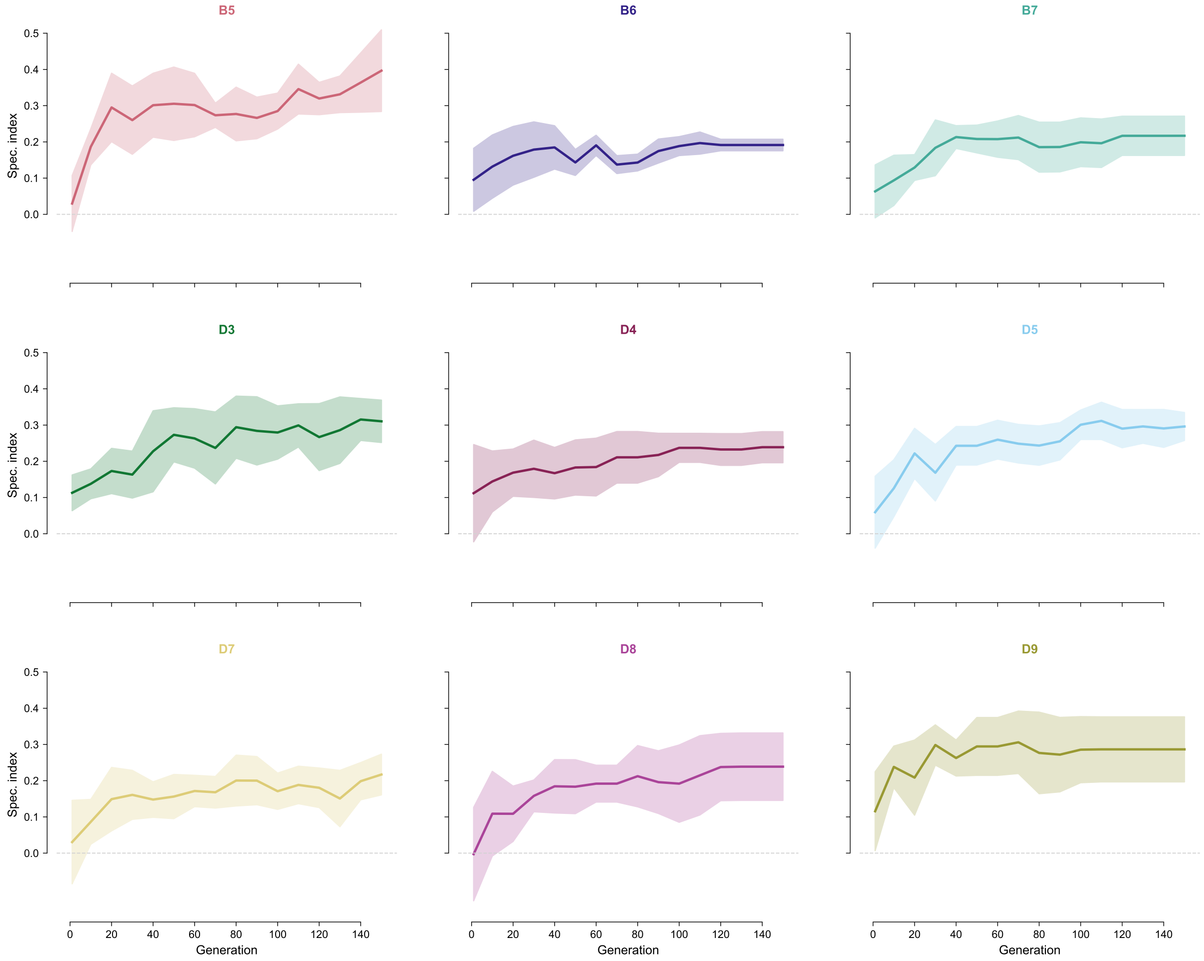


**Figure S22. Per-mouse specialisation index trajectories.** Specialisation index (mean ± SD, $n = 6$ replicates) over 150 generations for each of the 9 mice. Each panel corresponds to one mouse (colour-coded as in main figures). All mice reach positive specialisation indices; variance across replicates narrows with increasing generations.

Sensitivity commitment co-develops with behavioural specialisation. Figure S23 shows three temporal analyses from the same generation checkpoints used in the main specialisation trajectory. Panel A reproduces the population mean specialisation index for reference. Panel B shows mean per-neuron sensitivity variance over generations for specialists (blue) and generalists (orange) on a log scale; specialist variance remains elevated relative to the generalist level from approximately generation 20 onward, indicating that the higher sensitivity commitment seen at generation 150 builds progressively rather than emerging late. Panel C shows pairwise cosine similarity of sensitivity profile vectors: within the same mouse (solid) vs between mice (dashed). Within-mouse cosine similarity is marginally below the between-mouse baseline over training, but this small gap is not significant under a pseudoreplication-aware test (label-permutation $p = 0.076$; §2.6); we therefore read it only as the absence of a shared mouse-specific pathway signature, not as evidence that replicates commit to distinct pathways.

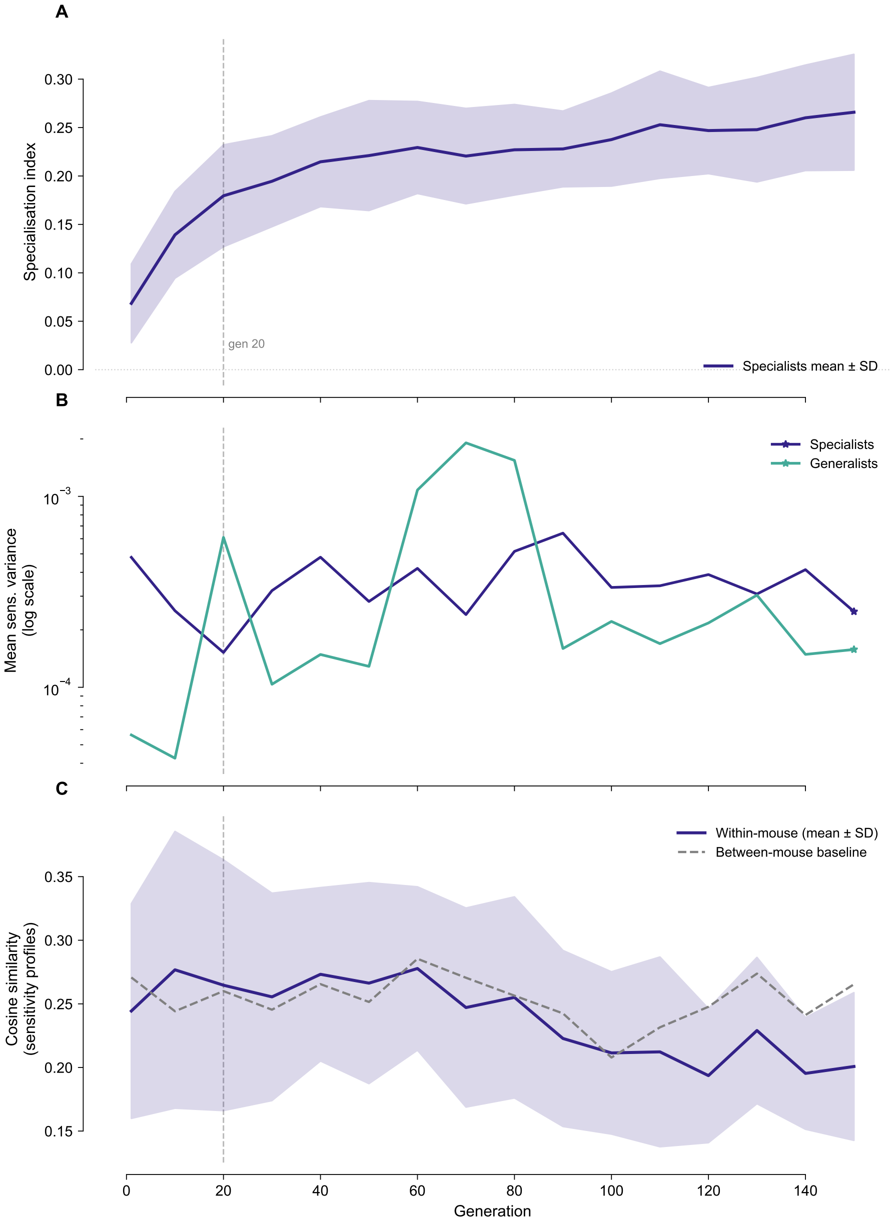


**Figure S23. Sensitivity commitment temporal co-development.** (A) Population mean behavioural specialisation index ± SD. (B) Mean per-neuron sensitivity variance over generations: specialists (blue) vs generalists (orange), log scale. Dashed vertical line marks generation 20. (C) Within-mouse pairwise cosine similarity of sensitivity profiles (solid line, ± SD shading) vs between-mouse baseline (dashed). Within-mouse similarity is marginally below the between-mouse baseline; this small gap is not significant under a pseudoreplication-aware test (label-permutation $p = 0.076$; §2.6), indicating no shared mouse-specific pathway signature rather than convergence to distinct per-replicate solutions.
